# Mechanism of receptor assembly via the pleiotropic adipokine Leptin

**DOI:** 10.1101/2022.12.01.518327

**Authors:** Alexandra Tsirigotaki, Ann Dansercoer, Koen H.G. Verschueren, Iva Marković, Christoph Pollmann, Maximillian Hafer, Jan Felix, Catherine Birck, Wouter Van Putte, Dominiek Catteeuw, Jan Tavernier, J. Fernando Bazan, Jacob Piehler, Savvas N. Savvides, Kenneth Verstraete

**Affiliations:** Unit for Structural Biology, Department of Biochemistry and Microbiology, Ghent University, Ghent, Belgium; Unit for Structural Biology, VIB-UGent Center for Inflammation Research, Ghent, Belgium; Department of Biology/Chemistry and Center of Cellular Nanoanalytics, Osnabrück University, Osnabrück, Germany; Integrated Structural Biology Platform, Centre for Integrative Biology (CBI), Institut de Génétique et de Biologie Moléculaire et Cellulaire (IGBMC), CNRS UMR 7104, INSERM U1258, University of Strasbourg, Illkirch, France; PUXANO, PXN BioLabs. BlueChem Incubator, Antwerp, Belgium; VIB-UGent Center for Medical Biotechnology, Ghent, Belgium; Department of Biomolecular Medicine, Ghent University, Ghent, Belgium; ħ Bioconsulting llc, Stillwater, MN, USA

## Abstract

The adipokine Leptin activates its type I cytokine receptor (LEP-R) in the hypothalamus to regulate body weight and exerts additional pleiotropic functions in immunity, fertility, and cancer. However, the structure and mechanism of Leptin-mediated LEP-R assemblies has remained unclear. Here, we show that Leptin:LEP-R assemblies adopt an unprecedented structure within the type I cytokine receptor family featuring 3:3 stoichiometry. We validate Leptin-induced trimerization of LEP-R in the plasma membrane of living cells via multicolor single molecule microscopy. In mediating such assemblies Leptin undergoes drastic restructuring that activates its site III for binding to the Ig-domain of an adjacent LEP-R molecule in the complex. These interactions are abolished by pathological mutations linked to obesity. Collectively, our study uncovers an evolutionarily conserved Leptin:LEP-R assembly as a new mechanistic blueprint for Leptin-mediated signaling in physiology and disease, including insights into how the lowly abundant signaling-competent isoforms of LEP-R can productively participate in signaling.

## Main

Leptin, a four-helix-bundle hormone ^1, 2^, activates LEP-R to convey information about the energy deposit levels in the white adipose tissue to the hypothalamus and also assumes pleiotropic roles in immunity, fertility, and cancer ^3–7^. There, LEP-Rb (**Fig. 1a**), the longest of the six LEP-R isoforms ^8, 9^ becomes activated by Leptin to transduce signaling via Janus kinase/Signal Transducer and Activator of Transcription (JAK/STAT) ^10, 11^ to balance energy expenditure and food intake. Dysregulation of Leptin-mediated STAT-3 signaling via LEP-Rb is linked to obesity ^2, 10, 12–14^. Contrary to the restricted expression of LEP-Rb, the STAT3-signaling deficient short LEP-R isoforms (LEP-Ra,c,d,f) carrying a truncated intracellular region beyond the box 1 motif are widely expressed ^9^ and may facilitate the transport of Leptin across the blood-brain barrier ^15–17^, while providing a source for soluble LEP-R via ectodomain shedding ^18^. The heavily glycosylated ^19^ extracellular domain (ECD) of LEP-R is common to all isoforms and can be classified in the gp130 cytokine receptor family, albeit with a unique architecture. LEP-R features a central combination of modules that comprising an Immunoglobulin-like domain (LEP-R_Ig_), which is essential for Leptin-induced receptor oligomerization, and a cytokine-receptor homology (CRH) 2 module (LEP-R_CRH2_) responsible for high affinity Leptin binding ^20–24^. Flanking the LEP-R_Ig-CRH2_ core at its N-terminus is a STAT3-signaling redundant ^20, 21^ CRH1 module (LEP-R_CRH1_) with an N-terminal extension (NTD) and at its C-terminus two Fibronectin III-like (FnIII) domains (LEP-R_FNIII_) proximal to the cell membrane (**Fig. 1a**). The latter are imperative for LEP-Rb-mediated STAT signaling ^20, 25^. By contrast, all other ‘tall’ gp130-class cytokine receptors feature ectodomain stalks with three FnIII domains, where the additional membrane-proximal domain (with respect to LEP-1. R) could possibly pack vertically as suggested from observed homo- or heteromeric 2:2 complexes of such cytokine receptors ^26^.

A plethora of mutagenesis data over the last two decades had led to a molecular mechanism whereby Leptin employs two distinct functional sites, termed sites II and III, to establish LEP-R signaling complexes at the cell surface ^3^. Furthermore, the receptor signaling assembly mediated by Leptin has been proposed to adopt a 2:2 stoichiometry or multiples thereof ^24^, based on structural paradigms derived from receptor complexes of IL-6 ^27^ and GCSF ^28^ and structural models obtained from negative-stain electron microscopy (EM) ^29^ and Small-angle X-ray Scattering (SAXS) ^22^, in line with the generally accepted dimeric mode of JAK activation ^30–34^. However, functional complementation of LEP-R variants deficient in JAK2-binding and phosphorylation suggested that Leptin:LEP-R assemblies adopting a higher-order of oligomerization are possible ^21^. Furthermore, minimal signaling repression of LEP-Rb by LEP-Ra ^35^ and evidence of heteromeric complexes comprising LEP-Rb and LEP-Ra at the cell surface ^36^ established that Leptin-driven LEP-R complexes do not need to comprise exclusively the signaling-competent LEP-Rb isoform.

**Figure 1.**
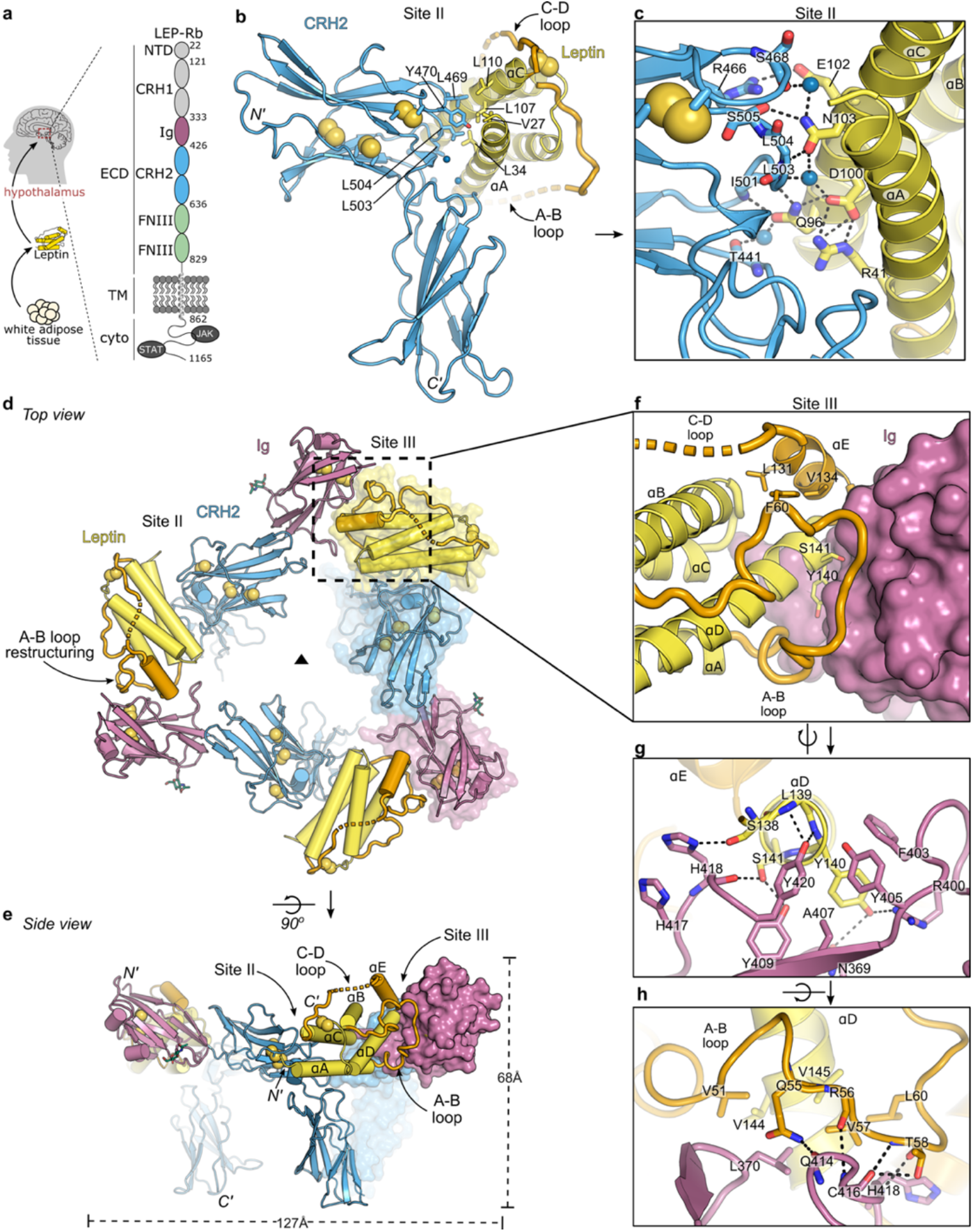
Structural basis of Leptin site II and site III binding to LEP-R to form assemblies with 3:3 stoichiometry. **a,** Domain organization of Leptin and the JAK/STAT signaling-competent long LEP-R isoform that is overexpressed in the hypothalamus. Leptin interacts with the LEP-R_CRH2_ (blue) and LEP-R_Ig_ (violet) domains to initiate signaling. **b,** Crystal structure of the glycan-trimmed mLeptin:mLEP-R_CRH2_ assembly featuring the site II interface. Interface residues involved in hydrophobic interactions are shown as sticks. Disulfide bonds are shown as yellow spheres. **c,** Focused view of the mLeptin:mLEP-R_CRH2_ site II interface featuring hydrogen bonds (dashed lines). Bridging water molecules are shown as blue spheres. **d-e,** Crystal structure of the glycan-trimmed mLeptin:mLEP-R_IgCRH2_ assembly from a top and side view. The asymmetric unit, highlighted as surface, comprises the high-affinity binary complex featuring the interaction site II. Adjacent copies of the binary complex assemble into a 3:3 complex mediated by interaction site III. **f,** The composite site III interface of Leptin with LEP-R_Ig_. Leptin docks on a surface cavity of LEP-R_Ig_ with the tip of α-helix D (blueprint Leptin site III residues in yellow sticks), and with the otherwise disordered A-B loop and α-helix E of the C-D loop that become ordered and internally stabilized with hydrophobic contacts (via residues shown as orange sticks). **g-h,** Site III interactions featuring Leptin’s α-helix D (panel g) and A-B loop (panel h) with LEP-R_Ig_.

We here present the structural and mechanistic basis for the assembly of Leptin-mediated LEP-R signaling complexes revealing an unprecedented assembly displaying 3:3 stoichiometry. These surprising findings provide the long-sought structural framework for further interrogation of Leptin-mediated signaling in physiology and disease, including how the lowly abundant signaling-competent isoforms of LEP-R can productively participate in signaling assemblies amid several highly abundant inactive LEP-R isoforms.

### Structural basis of high-affinity binding of Leptin to LEP-R via site II

The initial step towards LEP-R activation comprises Leptin binding with high affinity via a Leptin-specific site II-mediated interaction ^37^. To provide a structural snapshot of this critical interaction, we focused initially on complexes of Leptin with the minimal LEP-R fragment required for high-affinity Leptin binding ^20, 38, 39^ (**Extended Data Fig. 1a-c**), namely LEP-R_CRH2_. Ensuing crystal structures for glycan-trimmed mouse (m) and human (h) Leptin:LEP-R_CRH2_ complexes (**Extended Data Fig. 1a, Extended Data Table 1**) at 1.95 Å and 3.69 Å resolution, respectively, demonstrate that mouse and human complexes are akin (**Extended Data Fig. 1a**; root mean square deviation (r.m.s.d.) = 0.39Å, 161 Cα atoms), consistent with their high sequence identity (**Extended Data Fig. 1d**). Leptin utilizes αA and αC to bind the LEP-R_CRH2_ loop regions near the hinge of the CRH2 module (**Fig. 1b-c**), reminiscent of the site II interaction for homologous cytokines ^27, 28, 39, 40^. Notably, the loops connecting the αA-αB and αC-αD helices of both human and mouse Leptin are not resolved, in agreement with their dynamic nature in solution ^41^.

The crystal structure for the mLeptin:mLEP-R_CRH2_ assembly determined at high resolution provides a detailed description of this high-affinity cytokine-receptor interface. Both LEP-R_CRH2_ and Leptin employ evolutionary conserved residues to interact (**Extended Data Fig. 1d-e**) via hydrogen bonds and van der Waals contacts that bury approximately 1,600 Å^2^ of surface area (**Fig. 1b-c**, **Extended Data Fig. 4**). This interface is highly hydrated with six water molecules directly participating as bridging entities for hydrogen bonding (**Fig. 1b-c**, **Extended Data Fig. 4**).

Most interface contributions from LEP-R_CRH2_ stem from its first FNIII-like domain (**Fig. 1b**). At the heart of the interface, mLEP-R residues Leu469, Tyr470, Leu503 and Leu504 harbor at the hydrophobic junction of α-helices A-C of mLeptin (residues Val27, Leu34, Leu107, Leu110). Below the hydrophobic junction, hydrogen bonding dominates the interface, primarily via LEP-R_CRH2_ main chain – Leptin side chain contacts and/or bridging water molecules (**Fig. 1c**, **Extended Data Fig. 4**). mLeptin residues Arg41, Gln96 and Asn103 act as hydrogen bond hubs, with each participating in at least three different hydrogen bonds (**Fig. 1c**, **Extended Data Fig. 4**). The complex is stabilized by a salt bridge (mLeptin-Glu102 and mLEP-R-Arg466), supported by stacking of mLEP-R-Phe502 between mLEP-R-Arg466 and mLeptin-Asn99. The second FNIII domain of LEP-R_CRH2_ and the hinge interact exclusively with α-helix A of Leptin (**Fig. 1b, Extended Data Fig. 4**).

This observed site II corroborates previous structure-function studies of the Leptin:receptor interaction site II ^38–40, 42, 43^. In particular, mutations towards reduced affinity (mLeptin residues Asp30, Thr33, Lys36, Thr37, Arg41, Gln96, Asn103, Asp106, Leu107) ^39^, and impaired signaling (mLeptin residues Arg41 ^39, 43^, Gln96 ^39^ and Leu107 ^38^; mLEP-R_CRH2_ residues 501-504 ^38^), all map at the resolved site II interface (**Extended Data Fig. 4**).

### Leptin can assemble up to three LEP-R chains

Two long-standing questions in the activation mechanism of LEP-R by Leptin have been how Leptin mediates LEP-R oligomerization and what the precise stoichiometry of such complexes might be ^3^. Leptin-dependent oligomerization of LEP-R at the cell-surface was shown to be mediated by the Ig domain of LEP-R ^21^. To provide structural insights into this process, we initially focused on Leptin:LEP-R_IgCRH2_ complexes (**Extended Data Fig. 2**).

Minimally glycosylated Leptin:LEP-R_IgCRH2_ complexes were prepared for crystallographic studies and diffraction quality crystals were obtained for the mouse Leptin:LEP-R_IgCRH2_ (**Extended Data Table 1**). The crystallographic asymmetric unit of the resulting structure to 2.9 Å resolution in trigonal space group H32 contains one copy of the mLeptin:mLEP-R_IgCRH2_ complex that fully recapitulates the high-affinity complex seen in the mLeptin:mLEP-R_CRH2_ crystal structure (r.m.s.d= 0.4 Å for 252 aligned Cα atoms), with the mLEP-R_Ig_ domain emanating away from the LEP-R_CRH2_ module (**Extended Data Fig. 2a**). Application of the three-fold crystallographic symmetry generates a closed, ring-like Leptin:LEP-R_IgCRH2_ assembly with an unprecedented 3:3 stoichiometry (**Fig. 1d-e**) in the Type 1 cytokine receptor family ^44^. In this assembly, the LEP-R_CRH2_-bound Leptin exposes its site III epitope to bind the LEP-R_Ig_ domain from another Leptin:LEP-R_IgCRH2_ complex leading to a cross-over complex with 3-fold symmetry (C3).

In the LEP-R_IgCRH2_ fragment, LEP-R_Ig_ adopts a distinct configuration relative to the first FNIII-like domain of LEP-R_CRH2_, with a markedly outward shift compared to its IL6Rβ and GCSF receptor homologues (**Extended Data Fig. 3a**). This distinct geometry is mediated by the short linker at the LEP-R_Ig-CRH2_ boundary and stabilized by hydrophobic interactions at the interface between the Ig and CRH2 regions (**Extended Data Fig. 2a, S3a).** The interdomain configuration of the LEP-R_IgCRH2_ fragment, as well as the Leptin:LEP-R_CRH2_ and Leptin:LEP-R_Ig_ interaction modes agree well with structural models predicted by AlphaFold ^45, 46^, indicating that the experimentally determined assembly is supported by co-evolutionary relationships across the respective protein interfaces (**Extended Data Fig. 3c-f**).

**Figure 3.**
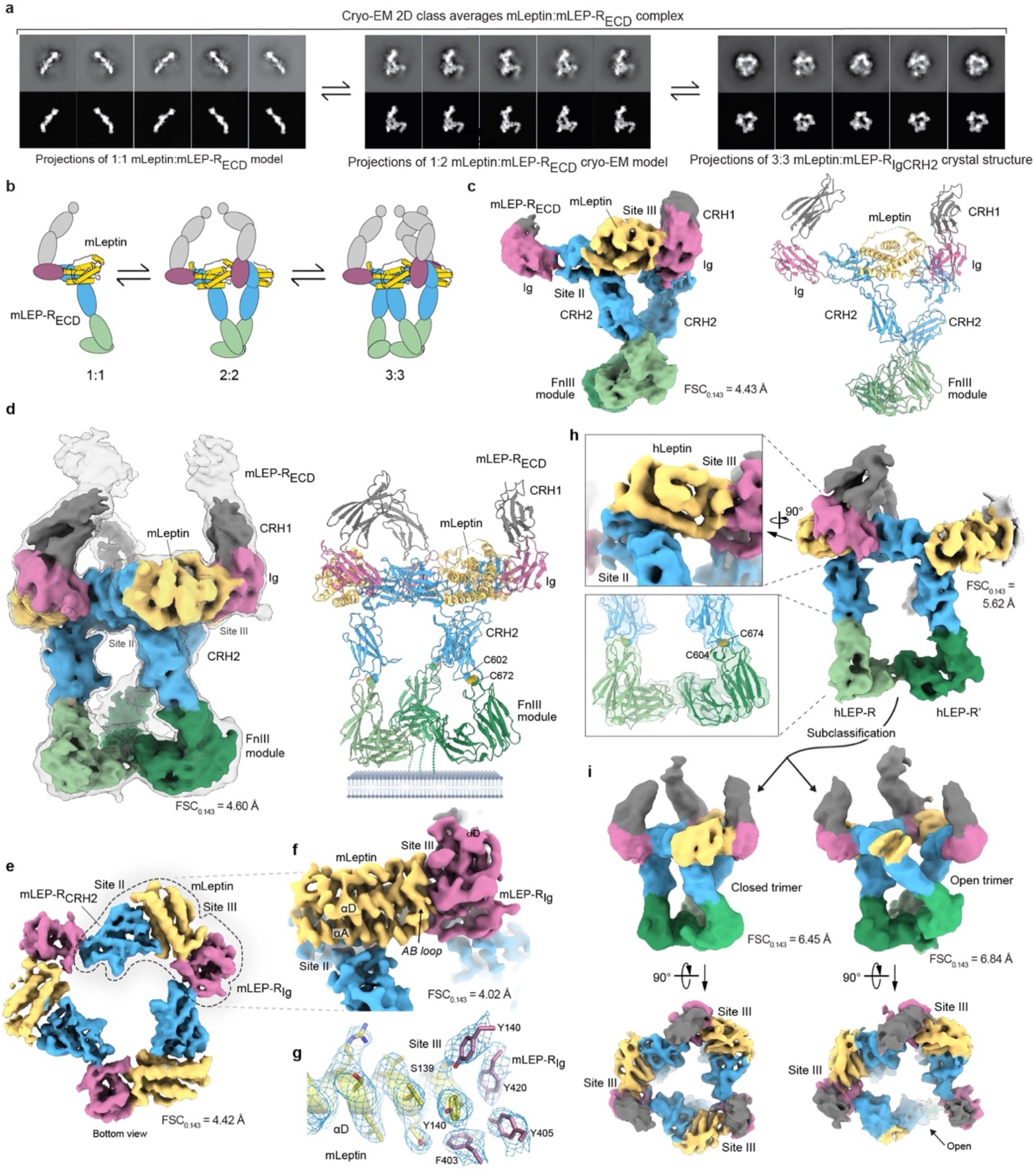
Structural characterization of the Leptin:LEP-R_ECD_ assembly via cryo-EM. **a,** Cryo-EM 2D class averages for the mLeptin:mLEP-R_ECD_ complex matched to projections of a 1:1 mLeptin:mLEP-R_ECD_ model obtained via AlphaFold (left), projections of the 1:2 mLeptin:mLEP-R_ECD_ cryo-EM model shown in panel c (middle) and projections of the determined 3:3 mLeptin:mLEP-R_IgCRH2_ crystal structure. **b,** Schematic representation of the mLeptin:mLEP-R_ECD_ assembly model. **c,** Cryo-EM map and real-space refined atomic model for the glycosylated 1:2 mLeptin:mLEP-R_ECD_ complex. The sharpened map is contoured at 0.39 V and colored per zone with mLeptin in yellow, the CRH1-module in grey, the Ig-domain in magenta, the CRH2 module in blue and the FnIII module in green. **d,** Cryo-EM maps and real-space refined atomic model for the glycosylated mLeptin:mLEP-R_ECD-tGCN4_ assembly. The sharpened map is contoured at 0.178 V. The non-sharpened map is shown as a transparent volume contoured at 0.09 V. **e,** Sharpened cryo-EM map for the mLeptin:mLEP-R_ECD-tGCN4_ assembly following local refinement around the mLeptin:mLEP-R_IgCRH2_ core region. The map is contoured at 0.313 V and displayed as a carved volume around mLeptin, the Ig-domain and the first domain of the CRH2-module with a carve radius of 4 Å. **f**, Sharpened cryo-EM map following symmetry expansion of the particle set in combination with local refinement around one IgCRH2:Leptin:Ig’ subcomplex. The map is contoured at 0.139 V. **g**, Sharpened map with DeepEMhancer following local refinement as in panel f. The map is centered around mLeptin-Y140 and contoured at 0.130 V and carved around the shown residues with a radius of 2.3 Å. **h,** Sharpened cryoEM map and real-space refined atomic model for the hLeptin:help-R_ECD-tGCN4_ complex contoured at 0.55 V. The map is colored via the same coloring scheme as for the mouse complex shown in panel d. The inset shows a cartoon representation of the atomic model and highlights the putative disulfide bridge in hLEP-R located at the interface between the CRH2 and the membrane-proximal FnIII-modules. **i**, Cryo-EM maps for the hLeptin:hLEP-R_ECD-tGCN4_ complex following further 3D-subclassification. The top panel shows the non-sharpened cryo-EM maps for reconstructions in closed and open state, contoured at 0.287 V and 0.245 V, respectively. The lower panel shows the corresponding sharpened maps in the closed and open states contoured at 0.45 V and 0.408 V.

Notably, in the crystal two trimeric Leptin:LEP-R_IgCRH2_ complexes are packed in a head-to-head orientation via reciprocal hydrophobic contacts of juxtaposing LEP-R_Ig_ domains (**Extended Data Fig. 2b**). In the absence of the N-terminal LEP-R domains, this head-to-head interface also stabilizes human and mouse Leptin:LEP-R_IgCRH2_ complexes in solution in an apparent 6:6 stoichiometry (**Extended Data Fig. 2c**), as determined by SEC-MALLS. These higher-order complexes are site III-dependent as the human Leptin variant that carries the ^141^ST^142^/AA (Leptin_a1_) mutations ^39^ was unable to form such complexes (**Extended Data Fig. 2c**), in agreement with the crystallographically-distilled 3:3 assembly, and are reduced to 3:3 assemblies in the context of the full-length LEP-R_ECD_ that contains the N-terminal LEP-R regions (**Fig. 2a**).

**Figure 2.**
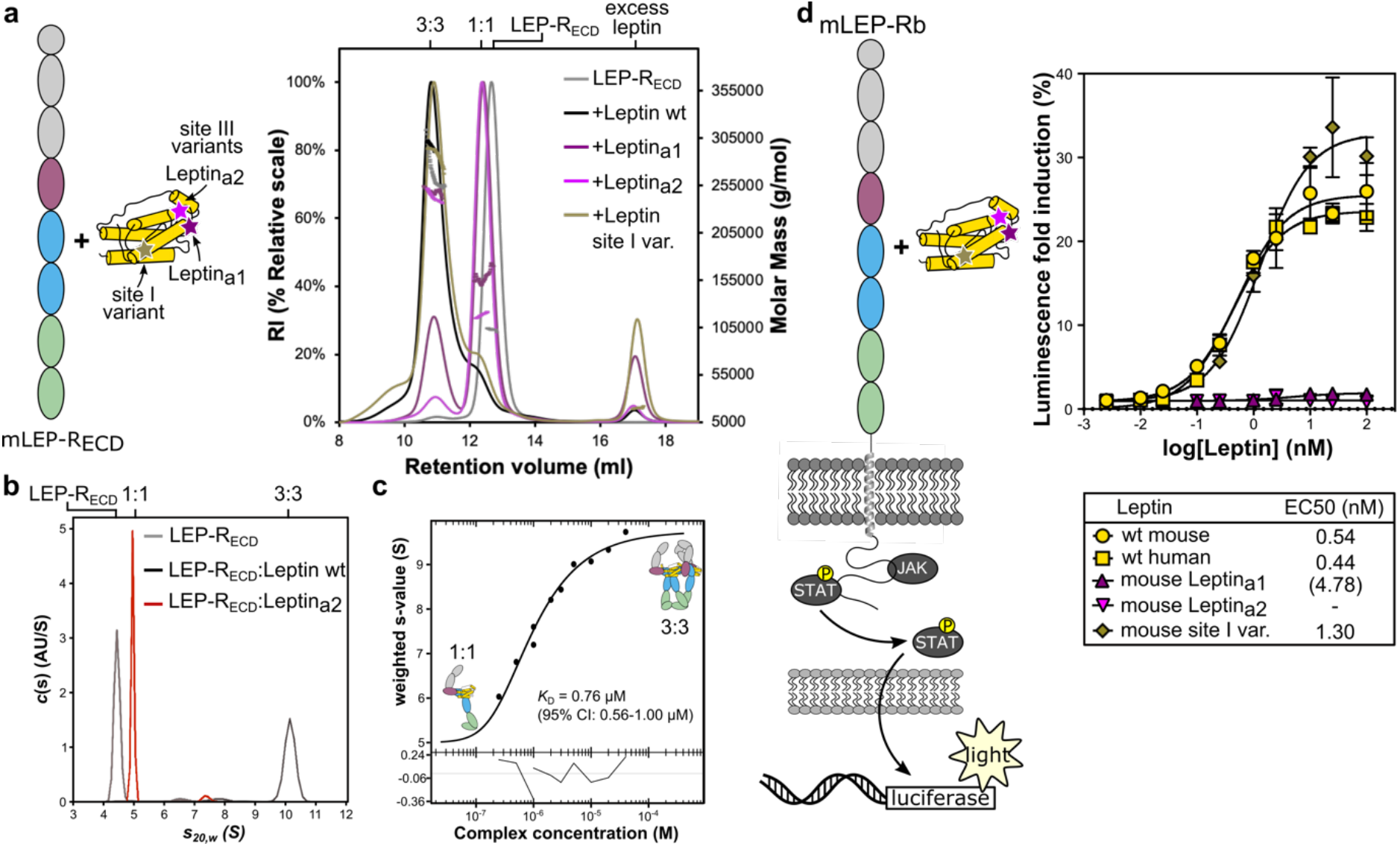
Leptin-dependent receptor oligomerization in solution and in living cells. **a,** Molar mass determination of glycan-trimmed mLEP-R_ECD_ and its complexes with Leptin variants by SEC-MALLS (Superdex 200 increase column). RI: Refractive index. **b,** Sedimentation coefficient distributions c(s) of glycan-trimmed mLEP-R_ECD_ alone or mixed at 1:1 molar ratio with mLeptin (wild type; wt) or mLeptina2 by AUC sedimentation velocity. **c,** Isotherm of the signal-weight-average s-values (*sw*) derived from the *c*(s) distributions presented in **Extended Data Fig. 6a**, and best-fit isotherm of a monomer-trimer self-association model with refined *KD* indicated. **d,** mLEP-Rb receptor activation by Leptin and variants probed by a STAT3 responsive luciferase reporter transiently transfected in HEK293T cells. Leptina1 only partially activates the receptor, in agreement with residual 3:3 oligomerization shown in panel a.

### Site III engagement evokes large conformational changes in Leptin

Leptin binds the Ig-domain of LEP-R with three structural elements contributed by residues located in the long A-B loop (residues 46 – 60), α-helix E (residues 130-137) located in the – D loop region and residues in the N-terminal region of α-helix D (residues 138-145) (**Fig. 1f-h**), Collectively, this composite interface leads to a total buried surface area of 1,600Å^2^ upon binding LEP-R_Ig_.

A comparison with the structure for hLeptin in the unbound state ^1^ and determined structures for the binary Leptin:LEP-R_CRH2_ complexes shows how the Leptin A-B and C-D loop regions restructure upon LEP-R_Ig_-binding (**Fig. 1b,d**, **Extended Data Fig. 2e**). Moreover, the N-terminal tip of α-helix D gains an extra turn upon LEP-R_Ig_-binding. The extended N-terminal tip of α-helix D contains the highly conserved residue Tyr140 (**Fig. 1f-g**, **Extended Data Fig. 1e**). Aromatic residues at this position comprise a blueprint of cytokine:Ig interactions in the gp130 family ^27, 28^ (**Extended Data Fig. 3b**). However, the Leptin:LEP-R_Ig_ interface is unique compared to its IL6:IL6Rβ and GCSF:GCSFR homologues. Instead of using a rather flat docking surface (**Extended Data Fig. 3b**), the LEP-R_Ig_ forms a surface cavity decorated with aromatic residues (mLEP-R-Phe403, -Tyr405, -Tyr409, -Tyr420) in parallel-displaced and T-shaped pi-stacking (**Fig. 1g**). This cavity embraces the tip of α-helix D and accommodates the aromatic ring of Tyr140 in the heart of the pocket, which contributes to T-shaped pi-stacking with mLEP-R-Phe403. The α-helix D is translated by 8.5Å on the z-axis relatively to IL6 and GCSF for shape complementarity with LEP-R_Ig_ (**Extended Data Fig. 3b**). In addition to T-stacking, the high tyrosine content in the LEP-R_Ig_ surface cavity mediates additional interactions via hydrogen bonding via hydroxyl groups (**Fig. 1g**). Strong hydrogen bonds (2.6Å distance) are formed between Ser141 at Leptin’s tip of α-helix D and mLEP-R-Tyr409 (hydroxyl group) and mLEP-R-His418 (carbonyl group) of LEP-R_Ig_, all ubiquitously conserved and important in signaling ^24, 39^. Moreover, this provides a structural rationale for the antagonistic ^141^ST^142^/AA (Leptin_a1_) variant ^39^ that carries mutations at the tip of α-helix D. Another signaling crucial residue, mLEP-R-Leu370 ^24^ of the β4 – β5 loop, packs against Val144-145 in α-helix D and Val51 and 57 of the A-B loop of Leptin forming a hydrophobic network (**Fig. 1h**).

Part of the C-D loop in Leptin becomes ordered in the LEP-R_Ig_-bound state, stabilizing an amphipathic two-turn α-helix, referred to as α-helix E in ref ^1^. This positions on top of the LEP-R_Ig_ cavity interacting with the β8 – β9 loop through van der Waals contacts and a hydrogen bond (mLeptin-Ser138 – mLEP-R-His418 of the His-His motif; **Fig. 1f,g**). Markedly, the A-B loop of Leptin is a major contributor to the interface, accounting for 53% of of Leptin’s site III interface area. It seals the interface by lining most of the rim of the LEP-R_Ig_ cavity, engaging in van der Waals contacts and hydrogen bonds (**Fig. 1f,h**, **Extended Data Fig. 4f**).

The hydrophobic packing against α-helices B-D (**Extended Data Fig. 2e**) stabilizes the Ig-binding competent conformation of the A-B and C-D loops. The αB-αD hydrophobic hub becomes occupied by the A-B loop, while α-helix E is stabilized in a V-configuration relatively to α-helix D. Residues ^62^FI^63^ bridge together the A-B loop, the four-helix-bundle and α-helix E. This rationalizes the antagonistic properties of Leptin variants affecting the binding conformation of site III to mediate the Leptin:LEP-R interaction interface, such as the ^62^FI^63^/AA Leptin variant ^47^, and the enhanced antagonistic nature of the _60_LDFI^63^/AAAA (**Fig. 2d**), which includes residue Leu60 directly binding LEP-R ^47^.

Collectively, the drastic restructuring of Leptin’s flexible A-B and C-D loops to enable LEP-R_Ig_ binding via site III is expected to be associated with an entropic penalty. This entropic cost might be compensated in part by the transient formation of α-helix E in the unbound state ^48^, and by the transient anchoring of the loops onto the hydrophobic hub formed by α-helices B-D (**Extended Data Fig. 2e**) ^41^, similarly to the crystallographic packing of human Leptin in the unbound state ^1^.

### Mapping of pathogenic mutations on the Leptin:receptor assembly

Several mutations in Leptin and LEP-R, including deletions, frameshifts and single nucleotide polymorphisms (SNPs), are associated with severe obesity and other diseases ^12, 49^. Here we focus on pathogenic SNPs with no currently known effect on secretion levels and stability (**Extended Data Table 2, Extended Data Fig. 5**).

The pathogenic LEP-R A409E (Ala407 in mouse) and Leptin S141C, V145E mutations localize at the site III interface (**Fig. 1g; Extended Data Fig. 5b**). In vitro studies on LEP-R A409E and Leptin V145E show that these mutants retain site II affinity but are signaling deficient (**Extended Data Table 2)**. The other two confirmed pathogenic mutations in Leptin, D100Y and N103K, map on the site II interface (**Fig. 1c; Extended Data Fig. 5c**), in agreement with their abolished LEP-R binding phenotype ^50–52^. The C604S mutation in human LEP-R does not map at the interface but would disturb the putative disulfide bridge between residues Cys604 and Cys674 (mLEP-R-Cys602, -Cys672; **Extended Data Fig. 5d,f**) that likely plays a key role to project the membrane-proximal FnIII-modules in a signaling competent conformation.

Thus, we corroborate pathological and functional interrogation mutations in Leptin and LEP-R, providing the missing consolidated resource for basic and clinical research.

### Site III interactions drive signaling Leptin:LEP-R assemblies

To investigate the stoichiometry of complexes of Leptin and variants thereof in complex with full-length LEP-R_ECD_, we employed SEC-MALLS and analytical ultracentrifugation (AUC) sedimentation velocity experiments (**Fig. 2a-c**, **Extended Data Fig. 6**). The biological activity of Leptin variants was interrogated in a STAT3 responsive luciferase reporter assay in HEK293T cells (**Fig. 2d**).

We found that the glycan-trimmed mLEP-R_ECD_ is monomeric in solution and forms 3:3 complexes with mLeptin (**Fig. 2a-b**), in a reversible and concentration-dependent manner (**Extended Data Fig. 6a, Fig. 2c**). This 1:1 to 3:3 transition is of low affinity in solution (*K*_D_ = 0.76 – 1.2 μM; **Fig. 2c**) and independent of the membrane proximal FNIII domains (**Extended Data Fig. 6c**). The formation of 3:3 complexes in solution and at the cell surface is site III-dependent as Leptin_a1/a2_ antagonists (^141^ST^142^/AA and ^60^LDFI^63^/AAAA respectively) bind to the receptor with a 1:1 stoichiometry but are impaired towards forming higher-order oligomers (**Fig. 2a,b**) and receptor activation (**Fig. 2d**). In addition, mLEP-R_ECD_ bearing the pathogenic A407E mutation at site III remains largely monomeric upon Leptin addition (**Extended Data Fig. 6b**). In contrast, mutating a previously proposed LEP-R binding site on Leptin towards a potential 2:4 Leptin:LEP-R assembly ^24^, termed ‘Site I’, located in the C-terminal region of α-helix D does not disturb the 3:3 assembly and this mutant also remains signaling competent (**Fig. 2a,d**). Spontaneous Leptin:LEP-R_ECD_ complexes in solution and signaling complexes are thus driven by the same interaction interfaces as in the crystallographically distilled assembly.

We also interrogated complex formation of glycosylated mLEP-R_ECD_ in solution using SEC-MALLS and found that the mLEP-R_ECD_ oligomerizes in a Leptin-dependent manner at similar concentrations as its glycan-trimmed counterpart, albeit with a lower – approximately 2:2 – apparent stoichiometry (**Extended Data Fig. 6d**). In contrast, glycosylation of the mLEP-R_ECD-_Δ_FNIII_ segment has no effect on the oligomerization towards a 3:3 mLeptin:mLEP-R_ECD-_Δ_FNIII_ complex in solution (**Extended Data Fig. 6c,e**). These data suggest that glycosylation at the membrane-proximal LEP-R_FNIII_ domains – which carries 5 potential N-linked glycosylation sites – may destabilize the 3:3 mLeptin:mLEP-R_ECD_ complex in solution, possibly via steric hindrance.

We were unable to demonstrate the formation of higher-order hLeptin:hLEP-R_ECD_ complexes with glycosylated hLEP-R_ECD_ in solution as only 1:1 hLeptin:hLEP-R_ECD_ complexes are observed via SEC-MALLS (**Extended Data Fig. 6f**). Glycan-trimmed hLEP-R_ECD_ precipitates upon Leptin addition, preventing further analysis. Strikingly, we found that although hLeptin does not support stable clustering in solution with hLEP-R_ECD_ and mLEP-R_ECD_, mLeptin forms 3:3 complexes in solution with glycosylated hLEP-R_ECD_ (**Extended Data Fig. 6g-h**). Efforts to stabilize hLeptin:hLEP-R complexes with 3:3 stoichiometry by “murinizing” hLeptin proved unsuccessful (**Extended Data Fig. 6i**). Nevertheless, hLeptin induces site III-dependent receptor activation in cells with similar EC_50_ values as mLeptin, and hLeptin and mLeptin exhibit full species cross-reactivity (**Fig. 2d, Extended Data Fig. 6k**). Moreover, site III-dependent activation appears significantly more efficient than activation by an agonistic anti-hLEP-R_ECD_ antibody ^53^ that dimerizes the human receptor in a Leptin-independent fashion (**Extended Data Fig. 6j,k**).

Collectively, these biophysical studies show that Leptin and the LEP-R ectodomain can form site III-dependent 3:3 complexes in solution with an affinity in the low micromolar range. At the membrane, the clustering of 1:1 Leptin:LEP-R complexes towards transmembrane higher-order signaling assemblies is expected to be facilitated by the reduced dimensionality of the membrane, which lowers the entropic cost of association of membrane-embedded receptors, as well as by possible additional contacts with the membrane, and interactions between the TM-helices and intracellular regions of adjacent LEP-R chains.

### Structure of the complete extracellular Leptin:LEP-R_ECD_ assembly

To obtain additional structural and mechanistic insights, the glycosylated Leptin:LEP-R_ECD_ assembly was characterized using single-particle cryo-EM analysis. Dilution of a concentrated mLeptin:mLEP-R_ECD_ complex sample just prior to cryo-EM grid preparation revealed distinct oligomeric states demonstrating that mLeptin:mLEP-R_ECD_ can assemble into complexes of up to 3:3 stoichiometry (**Fig. 3a,b**, **Extended Data Fig. 7a**), in line with our SEC-MALLS and AUC experiments. All three observed states appeared to populate preferred orientations, preventing analysis towards high-resolution 3D reconstruction. Anisotropic, cryo-EM reconstruction applied to an apparent intermediate complex (FSC_0.143_ resolution = 4.43 Å) revealed an open assembly in which mLeptin bridges two mLEP-R_ECD_ chains via sites II (LEP-R_CRH2_) and site III (Lep-R_’Ig_) interactions to elicit contacts between the membrane-proximal FnIII-modules of the two LEP-R chains (**Fig. 3c, Extended Data Fig. 7b**). Notably, the cryo-EM map of this complex lacks evidence for a second Leptin molecule at the CRH2’ site which may be linked to the apparent increased structural heterogeneity in this region and/or partial complex dissociation under these experimental conditions.

To facilitate cryo-EM analysis of the 3:3 mLeptin:mLEP-R_ECD_ complex, the complex was stabilized by fusion of the trimerizing GCN4 isoleucine-zipper motif ^54, 55^ (tGCN4) to the C-terminus of mLEP-R_ECD_ (**Extended Data Fig. 8a**). Non-stabilized glycan-trimmed mLeptin:mLEP-R_ECD_ complexes are structurally similar to the tGCN4-stabilized assembly in solution as probed by SAXS (**Extended Data Fig. 8b**), confirming that the tGCN4-fusion stabilizes spontaneously formed complexes.

Cryo-EM reconstructions in C1 symmetry to an FSC_0.143_ resolution of 4.60 Å for the glycosylated mLeptin:mLEP-R_ECD-tGCN4_ complex (**Extended Data Fig. 8c**) show an extended, pseudo-C3-symmetric assembly in which three Leptin:LEP-R_ECD_ complexes interact via site III mediated interfaces between Leptin and the Ig-domain of adjacent LEP-R chains as seen in the 3:3 Leptin:LEP-R_IgCRH2_ crystal structure (**Fig. 3d,e**, **Extended Data Fig. 10a,b**). To possibly resolve this pseudo C3-symmetric assembly at a higher resolution, we applied symmetry expansion in combination with local refinement. This approach drastically improved the map quality (FSC_0.143_ = 4.02 Å, **Fig. 3f,g** and **Extended Data Fig. 8d**) and demonstrates that the CRH2:Leptin:Ig’ subcomplexes form well-defined structural units within the ring-like core region of the complex.

The N-terminal mLEP-R_CRH1_ and membrane-proximal mLEP-R_FnIII_ modules emanate above and below this ring-like core region, respectively. The N-terminal domain of the LEP-R_CRH1_ modules is largely disordered in the cryo-EM maps. The membrane-proximal regions of the adjacent mLEP-R chains arrange in a non-symmetric manner whereby two receptor legs interact via the tips of their LEP-R_FnIII_ modules, and with the LEP-R_FnIII_ module of the 3rd receptor leg being less defined (**Extended Data Fig. 10c**). The membrane-proximal FnIII-module of mLEP-R_ECD_ adopts a bent, compact conformation as seen in gp130 ^56^. This bent conformation is also observed in a 1.75 Å crystal structure for mLEP-R_FnIII_ in complex with the anti-mLEP-R nanobody VHH-4.80 (**Extended Data Table 1, Extended Data Fig. 5e**), indicating that the two FnIII-like subdomains form a rigid structural unit, in part stabilized via evolutionary conserved Trp residues (Trp644, Trp783) across the interdomain interface ^56^.

Single-particle cryo-EM analysis for the corresponding hLeptin:hLEP-R_ECD-tGCN4_ complex (**Extended Data Fig. 9**) results in a reconstruction to an FSC_0.143_ resolution of 5.62 Å that shows how two hLeptin:hLEP-R_ECD_ subcomplexes form a site III-mediated assembly as seen in the cryo-EM analysis for the mLeptin:mLEP-R_ECD-tGCN4_ complex, while the density for a third Leptin:LEP-R subunit is not well-defined indicating structural heterogeneity (**Fig. 3h, Extended Data Fig. 10d,e**). Ensuing subclassification shows that the third hLeptin:LEP-R_ECD_ subcomplex can adopt two conformations towards closed and open trimeric assemblies (**Fig. 3i, Extended Data Fig. 9b**). This observation may be linked to the apparent weaker site III interaction for the human as compared to the mouse complex.

In conclusion, our cryo-EM analysis establishes that both mouse and human Leptin have the potential to oligomerize LEP-R via the crystallographically observed site II and site III interactions to elicit contacts between the membrane-proximal LEP-R_FnIII_ modules. The conformation of LEP-R_ECD_ as observed in the cryo-EM density corresponds remarkably well to the structural prediction via AlphaFold suggesting that the observed interdomain orientations in LEP-R_ECD_ agree with the co-evolutionary relationships between the structural domains in LEP-R_ECD_ (**Extended Data Fig. 10f**). In addition, the orientation of the membrane-proximal LEP-R_FnIII_ modules – which are required for LEP-Rb-mediated signaling ^20, 25^ – is likely restrained by a disulfide bridge between mLEP-R residues Cys602 and Cys672 across the CRH2 and FnIII interface (**Extended Data Fig. 5f**). The functional importance of this disulfide bridge to project the membrane-proximal FnIII-modules in a signaling competent conformation is underscored by the pathogenic C604S mutation in human LEP-R (**Extended Data Fig. 5a,d**).

We note that the structural plasticity observed in our cryo-EM samples might be exaggerated compared to the corresponding transmembrane complexes, as these would be expected to be further restrained by anchorage to the transmembrane helices and possible interactions between the LEP-R_FnIII_ modules and the outer plasma membrane region. It remains to be seen whether this plasticity is mechanistically linked to JAK/STAT activation.

Finally, in order to assess the evolutionary persistence of the observed human and mouse Leptin:LEP-R hexameric assemblies, we used AlphaFold-Multimer ^57, 58^ to derive 3:3 complexes for a series of Leptin:LEP-R complexes in animals, focusing on the jawed vertebrates where 4-helix-bundle cytokines and their receptors can be detected ^59^. The corresponding 3:3 Leptin:LEP-R_IgCRH2_ model complexes strongly converge on hexameric solutions that closely superpose with the crystallographically distilled 3:3 mLeptin:mLEP-R_IgCRH2_ assembly (**Extended Data Fig. 3g**). This suggests that Leptin conserves the ability to nucleate the assembly of 3:3 LEP-R ectodomain complexes from the very earliest Leptin signaling systems in jawed vertebrates.

### Leptin induces receptor trimerization at the cell surface

Receptor dimers are fundamentally required for JAK activation ^60, 61^ but higher order stoichiometry such as tetrameric receptors comprising two pairs of JAKs are also found in the class I and II receptor families. Recent experimental advances in multicolor single-molecule imaging have enabled to quantify receptor stoichiometries of signaling complexes in the plasma membrane of live cells ^62^. These techniques revealed that the subunits of type I/II cytokine receptors are monomeric in the resting state and associate into well-defined dimeric or tetrameric signaling complexes largely matching the complexes observed in *vitro* ^62–67^. Given the 3:3 Leptin-LEP-R stoichiometry that we consistently identified in *vitro*, we sought to investigate receptor assembly in the plasma membrane of living cells. To this end, we conducted triple-color single-molecule imaging by total internal reflection fluorescence microscopy (smTIRFM) ^62^ using N-terminally ALFA-tagged mLEP-Ra and mLEP-Rb labelled via nanobodies in three different colors (**Fig. 4a**).

**Figure 4.**
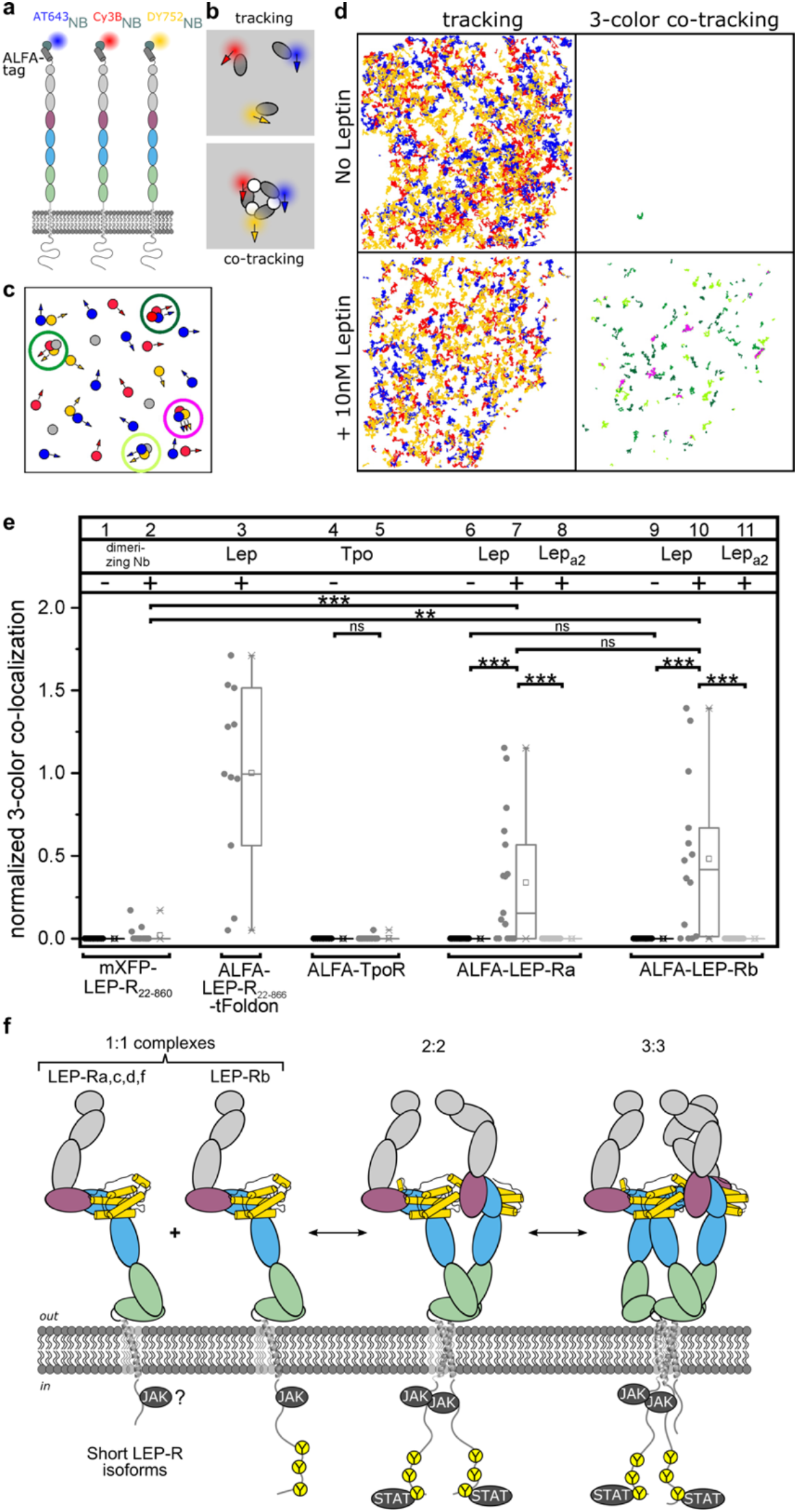
Leptin-induced assembly of LEP-R trimers at the cell surface. **a,** Quantifying receptor assembly in the plasma membrane of live cells by triple-color single molecule co-tracking. mLEP-R with N-terminal ALFA-tag at the surface of HeLa cells were stochastically labeled with the indicated dyes conjugated to anti-ALFA-tag nanobodies. **b**, Trimeric receptors are identified by triple-color co-localization and co-tracking analysis. **c**, Detection of trimers is limited by the random and incomplete receptor labeling as well as co-tracking fidelity. Therefore, relatively high levels of receptor dimers (green circles) are expected in addition to triple-color trimers (magenta circles). **d**, Representative triple-color co-tracking analysis of mLEP-Rb in the absence (top) and presence (bottom) of mLeptin. Single molecule trajectories (left) and co-trajectories (right) with dual- and triple-color co-trajectories identified in green and magenta, respectively. **e,** Relative number of triple-color co-trajectories observed for unstimulated and stimulated receptors, normalized to the mXFP-LEP-R_22-866-tFoldon_ stimulated with mLeptin. **f,** Potential Leptin-mediated homo- and heteromeric LEP-R assemblies at the cell surface.

Interactions between LEP-R subunits were quantified by dual- and triple-color single molecule co-tracking as exemplarily depicted in **Fig. 4b and c**. As expected, no co-diffusion of LEP-R was observed in the resting state, confirming receptor monomers in the absence of Leptin (**Fig. 4d,e**, **Extended Data Fig. 11a,b, Supplementary Movie 1**). However, upon ligand stimulation, substantial three-color (3C) co-diffusion was observed for LEP-R, but not TpoR, which we used as a control for a strictly homodimeric class I cytokine receptor ^65^ (**Fig. 4d,e**, **Extended Data Fig. 11a,b, Supplementary Movie 1**). Likewise, dimerization of LEP-R via an engineered cross-linker did not yield significant 3C co-diffusion (**Fig. 4e**), corroborating that the higher stoichiometry of Leptin-induced LEP-R complexes is based on ligand-mediated interactions.

To interpret the observed 3C co-diffusion, we engineered a constitutively trimerized mLEP-R variant by fusing a trimerization domain (“Foldon” from T4 fibritin) downstream of the transmembrane domain. Compared to the maximum 3C co-diffusion observed for this construct, we observed mean levels of ∼40% and 50% for mLeptin:mLEP-Ra and mLeptin:mLEP-Rb, respectively. These substantial 3C co-diffusion levels were accompanied by high dual-color (2C) co-diffusion (**Extended Data Fig. 11a, Supplementary Movie 1**), which can be explained by a stochastic labeling within trimers (**Fig. 4c**). Importantly, 2C and 3C co-diffusion were not detectable upon stimulation with Leptin_a2_, highlighting the key role of site III for the formation of LEP-R complexes in the plasma membrane. Changes in the diffusion properties upon stimulation with Leptin, however, were rather moderate, with the decrease in the diffusion constants and the increase in the immobile fraction being similar as observed for receptor dimerization (**Extended Data Fig. 11c,d**).

In conjunction with the relatively homogeneous fluorescence intensity of Leptin-induced LEP-R oligomers (**Supplementary Movie 1**), their largely uncompromised mobility highlights the lack of higher order clustering, in line with previous studies on other member of the class I/II cytokine receptor families ^62–67^. Rather, the co-tracking data strongly support our hypothesis that Leptin induces LEP-R trimers at the cell surface, which is also now illustrated structurally to near-atomic resolution by the plethora of structural and biophysical data we have presented. However, our observations are also in line with a mixture of LEP-R dimers and trimers at the cell surface.

## Discussion

After more than two decennia since the discovery of Leptin, we have elucidated here the structural basis of the Leptin:LEP-R assembly towards signaling, revealing a surprising architecture of circular hexamers. This establishes a unique cytokine-driven assembly mode within the Type I cytokine receptor family. This evolutionarily conserved Leptin:LEP-R assembly is mediated via the conserved and functionally validated sites II and IIII that spatially couple the tips of at least two membrane-proximal LEP-R_FNIII_ domains and depends on the dramatic restructuring on Leptin’s loop regions. The interaction of the LEP-R_FNIII_ domains is likely the key for triggering signaling by bringing the transmembrane helices, the intracellular region and consequently the associated JAK kinases to close proximity of each other for cross-activation ^32^. Corroborating our structural findings, we could confirm Leptin-induced formation of trimeric LEP-R complexes in the plasma membrane of living cells, rather than the canonical dimeric architecture of the family, which hitherto had served as the prevailing paradigm for LEP-R. However, the different mode of ligand engagement as compared to the paradigmatic receptors for Epo and growth hormone probably allows a more flexible stoichiometry at the cell membrane, that may also include open 2:2 Leptin:LEP-R complexes (**Fig. 4f**).

Considering the broad spectrum of LEP-R splice variants, the implications of such higher order and potentially variable receptor stoichiometries for intracellular JAK kinase activation and STAT3 signaling are intriguing. As the widely expressed ^9^ short LEP-R isoforms share an identical extracellular Leptin-binding region with LEP-Rb, these have the potential to form Leptin-dependent heteromeric complexes with LEP-Rb (**Fig. 4f**). Such Leptin-stabilized heteromeric LEP-R complexes at the cell surface have indeed been reported and may represent a major complex species at the cell surface ^36^. Although the short LEP-R isoforms contain the box 1 motif, the functional relevance of JAK recruitment by the short LEP-R isoforms ^9, 16, 68–70^ is unclear. Notably, LEP-Ra appears to show weak JAK2 binding and phosphorylation capacity ^71^, dependent on Box1. LEP-Ra is however unable to transactivate LEP-Rb towards STAT3 signaling in heterodimeric complexes as probed by chimeric LEP-Ra/b-IL5R receptors ^72^.

The minimal requirements for JAK/STAT signaling by LEP-R assemblies comprise two bound JAKs for cross-activation ^33, 73^ and at least one STAT3 docking site ^21^. As such, the Leptin:LEP-R system may have evolved to a higher-stoichiometry to facilitate LEP-Rb-mediated STAT3 signaling in the context of a mixture of long and short LEP-R isoforms. This is further supported by the weak repression of LEP-Rb signaling by LEP-Ra. Collectively, the structural insights reported here, which are supported by the large body of mutagenesis data on Leptin and LEP-R, provide a revised mechanistic blueprint to dissect Leptin-mediated signaling via the JAK-STAT axis in physiology and disease.

## ACKNOWLEDGEMENTS

We thank the staff of beamlines P12, P13 and P14 (Petra III, Deutsches Elektronen-Synchrotron), and Proxima 2A (SOLEIL) for technical support and beamtime allocation. We are grateful to the staff of the VIB-VUB facility for Bio Electron Cryogenic Microscopy (Brussels, Belgium) and the electron Bio-Imaging Centre (eBIC) at the Diamond Light Source (Didcot, UK) for their technical support. The pGL3-rPAPluc plasmid containing the luciferase gene was a kind gift of Dr. Frank Peelman (VIB-UGent, Ghent, Belgium). We thank Prof. dr. Jean-Pierre Timmermans from the Laboratory of Cell Biology and Histology (CBH) at Antwerp Centre for Advanced Microscopy (ACAM) for the use of the electron microscopy facilities for cryo-EM sample preparation.

This work was supported by grants from the Research Foundation – Flanders (grant G0G0619N to KV) and the VIB (to S.N.S.). This work benefited from access to the Integrated Structural Biology platform of the Strasbourg Instruct-ERIC centre IGBMC-CBI. Financial support was provided by FRISBI (ANR-10-INBS-0005) and Instruct-ERIC (PID 15107).

## AUTHOR CONTRIBUTIONS

A.T. prepared constructs, performed protein expression and purification with contributions from A.D. and K.V. A.T., K.H.G.V. and K.V. determined and analyzed crystallographic structures with contributions from S.N.S. K.V. collected and analyzed cryo-EM data, with contributions from J.F., W.V.P., and S.N.S. A.T. performed BLI, and SEC-MALLS experiments. SAXS data were analyzed by K.V. and A.T. I.M. performed cellular assays. M.H. and C.P. performed smTIRFM experiments and data analysis. J.F.B. carried out evolutionary structural analyses and J.P. supervised the smTIRFM experiments. C.B. performed SV–AUC experiments and analyzed data. D.C. and J.T. contributed critical reagents. A.T., K.V. and S.N.S. wrote the manuscript with contributions and approval from all authors. K.V. and S.N.S. conceived and supervised the project and procured funding.

## DECLARATION OF INTERESTS

The authors declare no competing interests.

## METHODS

### Materials availability

The HEK293S *MGAT1^-/-^* cell line and derivatives thereof cannot be freely distributed as some rights remain with the original authors.

Protein expression constructs generated in this study (as listed in **Supplementary Table 1**) are available via the BCCM/GeneCorner Plasmid Collection (http://bccm.belspo.be).

Crystallographic coordinates and structure factors have been deposited in the Protein Data Bank (PDB), and cryo-EM maps and models have been deposited in the Electron Microscopy Data Bank (EMDB) and PDB data banks with accession codes listed in **Tables S1** and S3.

### Protein expression in FreeStyle 293-F cells and purification from conditioned media

Human or murine LEP-R_ECD_, segments or variants thereof as well as human or murine Leptin and their variants (**Supplementary Table 1**) were produced in FreeStyle 293-F cells in suspension. Cells with density of approximately 1x10^6^ cells ml^-1^, grown in FreeStyle™ 293 Expression Medium (Gibco), were transfected with 1 μg ml^-1^ DNA using 2-fold excess of linear polyethylenimine (PEI; Polysciences), pre-incubated for 15 min in Optimem medium (Gibco). Transfected cell cultures were supplemented with 1% Penicillin-Streptomycin. Ex-Cell_®_ 293 Serum-Free Medium was added approximately 24 hours post-transfection to a final volume of 20%. Conditioned media containing the secreted recombinant protein were collected 4 days post transfection with centrifugation at 1,200 rpm for 15 min and were filtrated with PES filters (0.22 μm, Millipore).

His_6_-tag containing proteins were purified using metal ion affinity chromatography (IMAC; His-Trap FF, Cytiva). The column-captured material was washed sequentially with 1xPhosphate Buffer Saline (PBS; Sigma Aldrich), 1xPBS with 10mM Imidazole and eluted in 1xPBS supplemented with 250 mM Imidazole. The eluent was subjected to two rounds of SEC in 20 mM Hepes pH 7.4, 150 mM NaCl (HBS) using Superdex 200 increase 10/300 GL or Superdex 200 HiLoad 16/600 GL columns for LEP-R and Superdex 75 increase 10/300 GL for Leptin. For glycan-trimmed receptors, the IMAC-eluted material was concentrated to a maximal concentration of 1.5 mg ml_-_^1^ and incubated overnight with EndoH (1:50) at 4°C prior to SEC. For structural analysis (X-ray, SAXS) the His_6_-tag was removed with incubation with 1:40 Caspase-3 or TEV proteases, depending on the construct (**Auxiliary Supplementary Table 1**) in addition to EndoH. Protein samples were concentrated using centrifugal filters (Amicon). Leptin:LEP-R complexes were subsequently formed *in vitro* from the individually purified components at the desired concentrations. Exceptions comprise the mLeptin:mLEP-R_CRH2_ and hLeptin:hLEP-R_CRH2_ complexes used for crystallographic studies that were produced by co-transfection of Leptin and LEP-R_CRH2_ in 1:1 DNA ratio.

### Leptin expression in *E. coli* cells, refolding and purification from inclusion bodies

Expression of Leptin and variants thereof in *E. coli* facilitated high yield production of active, monomeric Leptin. BL21(DE3) cells were transformed with a pET15b vector encoding human or mouse Leptin, using Carbenicillin for selection. Cell cultures in LB (0.5 Lt; 37°C) were inoculated with overnight pre-cultures (1:100 back-dilution) and gene expression was induced at OD_600_=0.6 – 0.8 with 0.25 mM IPTG. Cells were harvested 3 to 5 hours post-induction by centrifugation (6,000 x g, 15 min, 4°C), resuspended in 50 mM Tris pH 8.0, 200 mM NaCl and lysed with a high-pressure homogenizer (Emulsiflex C3, Avestin). Inclusion bodies were harvested by centrifugation at 25,000 x g at 4°C for 20 min and were instantly frozen in liquid nitrogen. Inclusion bodies were resuspended in 6M Guanidine-HCl, 0.1M Tris pH 8.0 and homogenized with a Dounce homogenizer. The denatured sample was reduced with 5 mM DTT at 65°C for 10 min. Membranes and debris were removed by centrifugation at 20,000 rpm for 30 min at 4°C. The supernatant was refolded at 4°C with pre-chilled 0.1 M Tris pH 8.0, 10 mM reduced GSH, 1 mM oxidized GSSG mixture at 1 ml min^-1^ to a final Guanidine-HCl concentration of 0.1 M. Typically this also secured Leptin concentration of less than 0.1 mg ml^-1^ to minimize aggregation. Refolding was allowed overnight at 4°C with gentle stirring. Large aggregates were removed by centrifugation at 4,000 x g at 4°C for 15min and filtration (0.22 μm, Millipore). Refolded Leptin was then purified with IMAC (His-Trap FF, Cytiva) at 4°C, pre-equilibrated with 0.1 M Tris pH 8.0. Two washing steps were performed with 0.1 M Tris pH 8.0 and 0.1 M Tris pH 8.0, 10 mM Imidazole respectively prior to elution in 0.1M Tris pH 8.0, 250mM Imidazole. The eluent was concentrated and injected onto a HiLoad 16/600 Superdex 75 (Cytiva) column pre-equilibrated in HBS. Peak fractions corresponding to the monomeric refolded Leptin were polished with an additional SEC (HiLoad Superdex 75 or Superdex 75 increase).

### Preparation of protein complexes for crystallization

#### hLeptin: hLEP-R_CRH2_ complex

A codon-optimized cDNA fragment encoding the hLEP-R_CRH2_ receptor fragment (Uniprot P48357, residues 428-635) carrying the N516Q and C604S mutations was cloned into the pCAGGS vector in frame with the mouse Igh secretion signal (Uniprot Q32NZ5, residues 1-19) and an N-terminal TEV-cleavable hexahistidine-tag. A codon-optimized cDNA fragment encoding hLeptin (residues 22-167) was cloned in the pTwist-CMV-BetaGlobin vector (Twist BioScience) in frame with the mouse Igh secretion signal (Uniprot Q32NZ5, residues 1-19) and an N-terminal TEV-cleavable hexahistidine-tag. hLEP-R_CRH2_-N516Q/C604S and hLeptin were co-expressed in FreeStyle 293-F suspension cultures via PEI-mediated transient transfection in the presence of 20 μM kifunensine (Dextra Laboratories). The hLeptin: hLEP-R_CRH2_ complex was purified from the clarified conditioned medium via IMAC and SEC. N-linked glycans were trimmed by overnight incubation with EndoH using a 1:50 ratio (w/w) followed by SEC as a final polishing step using a Superdex 75 Increase column with HBS pH 7.4 as running buffer. The hLeptin:hLEP-R_CRH2_ complex (6 mg/mL) was subjected to crystallization trials using a Mosquito liquid handling robot (TTP Labtech) in sitting drop format. Crystals were obtained in condition F1 of the Index screen HT screen (0.2 M L-proline, 0.1 M HEPES pH 7.5, 10% PEG-3350) and cryo-protected with mother liquor supplemented with 20% PEG-400.

#### mLeptin: mLEP-R_CRH2_ complex

A codon-optimized cDNA fragment encoding the mLEP-RCRH2 receptor fragment (Uniprot P48356, residues 426 – 633) carrying the N514Q and C602S mutations was cloned into the pCAGGS vector in frame with the mouse Igh secretion signal and an N-terminal TEV-cleavable hexahistidine-tag. A codon-optimized cDNA fragment encoding mLeptin (Uniprot P41160, residues 22 – 167) was cloned in the pTwist-CMV-BetaGlobin vector in frame with the mouse Igh secretion signal and an N-terminal hexahistidine-tag. mLEP-R_CRH2_-N514Q/C602S and mLeptin were co-expressed in FreeStyle 293-F suspension cultures via PEI-mediated transient transfection in the presence of 20 μM kifunensine (Dextra Laboratories). The mLeptin: mLEP-R_CRH2_ complex was purified from the clarified conditioned medium via IMAC and SEC. The complex was incubated overnight incubation at room temperature with TEV protease and EndoH (1:50 and 1:50 w/w respectively) for cleavage of the His-tag and trimming of the N-linked glycans, respectively, followed by SEC as a final polishing step using a Superdex 75 Increase column with HBS pH 7.4 as running buffer. The mLeptin:mLEP-R_CRH2_ complex was concentrated to 7.6 mg mL^-1^ and subjected to crystallization trials using a Mosquito liquid handling robot (TTP Labtech) in sitting drop format. Crystals were obtained in condition H1 of the BCS screen HT screen (0.04 M calcium chloride dihydrate, 0.04 M sodium formate, 0.1 M Tris pH 8.0, 25% PEG Smear Low). Crystals were cryo-protected with mother liquor supplemented with 20% PEG-400.

#### mLeptin:mLEP-R_IgCRH2_ complex

For the mLeptin:mLEP-R_IgCRH2_ crystal structure, the mLEP-R_IgCRH2_ receptor fragment (Uniprot P48356 residues 328-633; C602S) was expressed in FreeStyle 293-F suspension cultures supplemented with the Mannosidase I inhibitor Kifunensine (Dextra Laboratories) at a final concentration of 20 μM. mLEP-R_IgCRH2_ secreted to the conditioned medium was purified using IMAC and was incubated with TEV and EndoH (1:20 and 1:50 w/w respectively) for cleavage of the purification tag and trimming of the N-glycans, respectively. The efficiency of the digestions was validated by SDS-PAGE and Western blot analysis using the 6X His Tag Monoclonal Antibody DyLight_TM_ 800 (Rockland). The receptor was incubated with 2-fold molar excess of tag-less mLeptin, refolded from *E. coli* inclusion bodies as described in *Vernooy et al., 2010* (ref^74^). The complex was concentrated to 6.1 mg ml^-1^ and injected onto a Superdex 200 increase in HBS. Fractions of the main peak, corresponding to a 6:6 complex as determined previously by SEC-MALLS, were concentrated to 4 mg ml^-1^ and subjected to crystallization trials using a Mosquito liquid handling robot (TTP Labtech) in sitting drop format with 100 nL protein mixed with 100 nL mother liquor in SwissSci 96-well triple drop plates and incubation at 20°C. Crystals were obtained overnight at 3.0 M Sodium formate, 0.1 M Tris pH 7.5 and at 1.0 M Ammonium sulfate, 0.1 M MES pH 6.5 of the ProPlex HT-96 screen (Molecular Dimensions). Crystals were cryoprotected in 4 M sodium formate + 10% (v/v) ethylene glycol prior to being cryocooled in liquid nitrogen.

#### mLEP-R_FnIII_:VHH 4.80 complex

A codon-optimized cDNA fragment encoding the mLEP-R_FnIII_ receptor fragment (Uniprot ID P48356 residues 633 – 827) and carrying the C672S and N668Q/N698Q/N726Q mutations was cloned in the pHLsec vector in frame with the vector’s secretion signal (Aricescu et al., 2005). A stop codon was inserted before the vector’s C-terminal His-tag. A cDNA fragment encoding the anti-mLEP-R_FnIII_ nanobody VHH-4.80 (ref. ^75^) was cloned also cloned in the pHLsec vector in frame with the vector’s secretion signal and in frame with the C-terminal His-tag. mLEP-R_FnIII_ and VHH-4.80 were co-expressed in HEK293S MGAT1_-/-_ suspension cultures via PEI-mediated transient transfection. The mLEP-R_FnIII_:VHH-4.80 complex was purified from the clarified conditioned medium via IMAC and following buffer exchange to HBS, the complex was incubated overnight at room temperature with EndoH using a protein target:EndoH ration of 1:30 (w/w). SEC was performed as a final polishing step using a Superdex 75 Increase column with HBS pH 7.4 as running buffer. The glycan-minimized mLEP-R_FnIII_:VHH-4.80 complex was concentrated to *2.5* mg ml^-1^ and subjected to crystallization trials using a Mosquito crystallization robot in sitting drop format with 200 nL protein mixed with 100 nL mother liquor in SwissSci 96-well triple drop plates and incubation at 20°C. Crystals were grown in the condition C4 of the BCS HT screen (0.2 M ammonium sulphate, 0.1 M sodium cacodylate pH 5.6 and 30% PEG Smear Medium) and cryoprotected with mother liquor supplemented with 30% (v/v) PEG-400.

### Crystallographic structure determination

Diffraction data was collected at the P13 and P14 (PETRAIII, DESY, Hamburg, Germany) and Proxima2A (SOLEIL, Paris, France) beam lines. All data were integrated and scaled using XDS ^76^ and AIMLESS ^77^ and data quality was analyzed by phenix.xtriage ^78^. Phases for the minimal Leptin:LEP-R_CRH2_ complexes were obtained by maximum-likelihood molecular replacement with PHASER ^79^ using the coordinates of human Leptin (pdb 1ax8) and the human LEP-R_CRH2_ fragment (pdb 3v6o). The determined structure of mLeptin:mLEP-R_CRH2_ was used as search model to solve the mLeptin:mLEP-R_IgCRH2_ complex structure. To solve the structure of the mLEP-R_FnIII_:VHH-4.80 complex search models were prepared based on pdb 1OHQ and a structural predicted model for mLEP-R via AlphaFold (AF-P48356-F1-model_v1). Model (re)building was performed in Coot ^80^ and refinement of coordinates and atomic displacement parameters was performed in PHENIX ^78^ and autoBUSTER ^81^. Model and map validation tools in Coot and the PHENIX suite were used throughout the work flow to guide improvement and to validate the quality of crystallographic models.

### Multi-Angle Laser Light Scattering (SEC-MALLS)

Protein samples of 100 μl were injected at the indicated concentrations onto a Superdex 200 increase 10/300 GL or a Superose 6 increase 10/300 GL column (Cytiva) pre-equilibrated in 20mM Hepes pH 7.4, 150 mM NaCl at a flow rate of 0.75 ml min_-1_. The column was coupled online with a UV-detector (Shimadzu), a miniDawnTReos (Wyatt) Multi-angle laser light scattering (MALLS) detector and an Optilab T-Rex (Wyatt) refractometer. The refractive increment value (dn/dc) was 0.185 ml g_-1_. Data was processed with the Astra 6.1.6 software. Band broadening was corrected for using reference measurements of BSA (Pierce). Glycosylated receptor complexes were subsequently subjected to protein conjugate analysis using a modifier refractive increment value (dn/dc) of 0.165 ml g_-1_.

### Biolayer Interferometry (BLI)

All measurements of binding kinetics and dissociation constants were performed using an Octet Red 96 (Forté Bio) in kinetics buffer (PBS, 0.1% (w/v) BSA, 0.02% (v/v) Tween-20) at 25°C. mLEP-R_CRH2_ or mLeptin, as indicated, were *in vitro* biotinylated using the EZ-Link™ NHS-PEG4-Biotinylation kit (Pierce) and desalted for removal of free biotin prior to immobilization on streptavidin-coated biosensors (Forté Bio) at 1.2 and 1.5 nm level respectively. The corresponding ligands, mLeptin and mLEP-R_ECD_ respectively, were prepared in two-fold dilution series in kinetics buffer. Parallel referencing with non-functionalized biosensors into all ligand concentrations as well as into kinetics buffers was employed. Data analysis was performed using Octet Analysis Studio 12.2 (Forté Bio) and binding curves were exported to Excel (Microsoft) for plotting of curves.

### Sedimentation velocity Analytical Ultracentrifugation (AUC)

The day prior to sedimentation velocity experiments the protein samples stored at -80°C were thawed and aggregates were removed by SEC (Superdex 200 10/30 GL and Superdex 75 10/30 GL for LEP-R and Leptin samples, respectively), equilibrated in SEC buffer (20 mM HEPES pH 7.4, 150 mM NaCl). SV–AUC experiments were conducted in a ProteomeLab XL-I analytical ultracentrifuge (Beckman Coulter) at 25 °C. The samples were loaded into AUC cell assemblies with 12- or 3-mm charcoal-filled Epon double-sector centerpieces. The sample cells were loaded into an 8-hole An-50 Ti rotor for temperature equilibration for 2–3 h, followed by acceleration to full speed at 42,000 rpm. Absorbance data at 290 nm (for concentrations at 40 µM), 280 nm (for concentrations in the range 1 to 20 µM) or 233 nm (for concentrations in the range 0.25 to 1 µM) were collected at 5-minute intervals for 15 h. The partial specific volumes of the proteins, buffer density and viscosity were calculated using the software SEDNTERP. Sedimentation data were time-corrected and modelled with diffusion-deconvoluted sedimentation coefficient distributions c(s) in SEDFIT 16.1c, with signal-average frictional ratio and meniscus position refined with nonlinear regression ^82^. Maximum entropy regularization was applied at a confidence level of 68%. Sedimentation coefficient distributions were corrected to standard conditions (water at 20°C, s20,W). Weight-averaged s-values (sw) at each concentration were determined by integrating c(s) distributions over the s-range of 4 to 12 S. Constructed sw isotherm were fitted with a monomer-trimer self-association model using SEDPHAT 15.2b to calculate the dissociation constant ^83^. The error in KD was estimated using the “Critical chi square for error surface projections” option in SEDPHAT with a confidence level of 95%. All plots were created in GUSSI ^84^.

### Leptin biological activity assay

HEK293T cells were transiently transfected with the mLEP-R_ALFA-tag_ and hLEP-R_ALFA-tag_ described in **Auxiliary Supplementary Table 1** and the pGL3-rPAPluc reporter construct containing the luciferase gene under control of the STAT3-inducible rat pancreatitis-associated protein-1 promoter ^85^. HEK293T cells were seeded into a 6-well plate and cultured overnight in DMEM (Invitrogen) and 10% fetal bovine serum (GibCo) in 5% CO2 at 37 °C. Next day, cells were transfected with mLEP-R_ALFA-tag_ or hLEP-R_ALFA-tag_ and the (optional: STAT3-responsive luciferase reporter construct) pGL3-rPAPluc DNA construct using linear polyethylenimine. After an overnight incubation, cells were detached using cell dissociation agent (Invitrogen) and 2% of the cell suspension was distributed per well of a 96-well plate. Cells were stimulated for 18h with increasing concentrations of human or mouse Leptin and mutant variants respectively. Afterwards, cells were lysed and the luciferase activity was measured by chemiluminescence in a Glomax luminometer (Promega) using luciferase substrate as previously described in *Peelman et al., 2004* (ref ^39^). Relative luciferase induction represents the stimulation factor normalized by the activity obtained with non-stimulated cells. Mean values were calculated from triplicates measured in parallel. Error bars represent standard deviations. The data were fitted to a log agonist versus response curve using GraphPad Prism.

### Single Molecule Total Internal Reflection Fluorescence Microscopy (smTIRFM)

For cell surface labelling, receptors were N-terminally fused to suitable tags using a pSems vector including the signal sequence of Igκ (pSems-leader). TpoR, mLEP-Ra, mLEP-Rb and mLEP-R_22-866_-tFoldon were fused to the ALFA-tag ^86^; mLEP-R_22-866_ was fused to non-fluorescent monomeric GFP (mXFP) ^65^. HeLa cells (ACC 57, DSMZ Germany) were cultured and transiently transfected as previously described ^87^. Anti-GFP and anti-ALFAtag nanobodies site specifically labelled with Cy3b, ATTO 643 and DY-752 via a C-terminal cysteine were added at concentrations of 1.5 nM each, at least 10 min before imaging.

Single-molecule imaging was carried out by triple-color total internal reflection fluorescence microscopy using an inverted microscope (IX-83, Olympus) equipped with a spectral image splitter (Multisplit V2, CAIRN) and a back-illuminated sCMOs camera (Hamamatsu ORCA Fusion-BT). Fluorophores were excited by simultaneous illumination with a 561 nm laser (MPB Communications), a 642 nm laser (MBP Communications) and a 730 nm laser (LuxX 730-50, Omicron). Image stacks of 150 frames were recorded for each cell at a time resolution of 33 ms per frame. Ligands (10 nM Leptin, 10 nM Leptin_a2_, 10 nM Tpo) and crosslinker nanobody (crosslinked LaG16) were incubated for 10 min before imaging. All imaging experiments were carried out at 23°C.

Triple-color time-lapse images were evaluated using an in-house developed MATLAB software (SLIMfast4C, https://zenodo.org/record/5712332) as previously described in detail ^62^. After channel registration based on calibration with fiducial markers, molecules were localized using the multi-target tracking algorithm ^88^. Immobile emitters were filtered out by spatiotemporal cluster analysis ^89^. For co-tracking, frame-by-frame co-localization within a cut-off radius of 100 nm was applied followed by tracking of co-localized emitters using the utrack algorithm ^90^. Molecules co-diffusing for 10 frames or more were identified as co-localized. Relative levels of co-localization were determined based on the fraction of co-localized particles ^91^. Diffusion properties were determined from pooled single trajectory using mean squared displacement analysis for all trajectories with a lifetime greater than 10 frames. Diffusion constants were determined from the mean squared displacement by linear regression.

### Small Angle X-ray Scattering (SAXS)

For mLeptin:mLEP-R_ECD_, the IMAC and SEC-purified glycan-trimmed receptor fragment was incubated with 1.5-fold molar excess of TEV-digested mLeptin refolded from *E. coli* inclusion bodies and injected onto a Superdex 200 increase 10/300 GL column in HBS, to remove the excess of Leptin. Samples of the complex were then prepared in HBS in a concentration range of 0.5 to 40 µM and were flash-frozen in liquid nitrogen. Samples in the same concentration range were also prepared for the mLEP-R_ECD_ alone, starting from the same glycan-trimmed receptor fragment sample used for the complex.

For mLeptin:mLEP-R_ECD-tGCN4_, His_6_-mLeptin refolded form *E. coli* inclusion bodies was used as a bait to pull down the tag-less mLEP-R_ECD-tGCN4_ from conditioned media of FreeStyle 293-F cultures supplemented with 20 µM Kifunensine using IMAC. The complex was subjected to two rounds of SEC (Superose 6 increase 10/300 GL) in HBS, with TEV and EndoH-digestion implemented before the last SEC. 1.5-fold molar excess of TEV-digested mLeptin was added to the complex prior to the last SEC to ensure full receptor occupancy. Samples of the complex were then prepared in HBS at 14 and 30 µM and were flash-frozen in liquid nitrogen. Prior to analysis, all samples were centrifuged at 13,000xg for 10min at 20°C.

In-solution SAXS data were measured on the P12 beamline of EMBL at the Petra III storage ring (DESY, Hamburg) in batch mode in HBS (20 mM HEPES, 150 mM NaCl, pH 7.2) at 25°C. The scattering data was collected in continuous flow mode using an automatic sample changer with 0.1 s exposure time per frame using 35 µL of sample and a flow rate of 10 µL/s resulting in 35 x 0.1s exposures. Scattering data was radially averaged and buffer subtracted using the automated SASFLOW pipeline^92^. Analysis of the radius of gyration and molecular weight were performed using Primus in the ATSAS suite ^93^.

### Cryo-EM data collection, image processing, structure modeling and refinement

#### mLeptin:mLEP-R_ECD_ complex

Prior to complex formation the N-terminal His-tag of refolded mouse Leptin was removed by an overnight TEV protease digest at room temperature using a target protein:protease ratio (w/w) of 50:1 followed by a polishing step via a Superdex 75 Increase SEC run. A 1.5 molar excess of mouse Leptin was then added to purified, N-terminally His-tagged mLEP-R_ECD_ produced from HEK293 FreeStyle cells in the presence of 20 μM kifunensine as described above, and the resulting complex was isolated via SEC using a HiLoad 16/600 Superdex 200 column with HBS as running buffer. Fractions corresponding to the mLeptin:mLEP-R_ECD_ complex were pooled, concentrated to 5 mg/mL, aliquoted, flash-frozen into liquid nitrogen and stored at -80 °C until further use. Immediately prior to cryo-EM grid preparation the mLeptin:mLEP-R_ECD_ complex was thawed and diluted to 0.2 mg/mL with HBS buffer and 4 μL sample was applied to a glow-discharged Quantifoil R 0.6/1 300 mesh golden grid coated with graphene (PUXANO), blotted for 1 s (blot force = 1) under 100% humidity at 22 °C and plunged into liquid ethane using an FEI Vitribot Mark IV. Grids were screened using an JEOL 1400 Plus (BECM, Brussels). Data collection was performed using a JEOL cryoARM 300 microscope equipped with a 6k X 4k GATAN K3 detector resulting in 7100 movies with a raw pixel size of 0.755 Å. Movies were motion corrected via MotionCor2 1.4.0 (ref. ^94^) and had their contrast transfer functions (CTFs) determined via patch-based CTF estimation as implemented in cryoSPARC v3.3.2 (ref. ^95^). Particles were extracted with a box size of 440 pixels with 2x binning. Initial high-resolution 2D classes were obtained via the blob picker function and reference-free 2D classification in cryoSPARC, These 2D classes were then used to seed template-based and neural network-based particle picking via Topaz 0.2.4 (ref. ^96^). Junk particles were removed by multiple rounds of 2D classification. High-resolution 2D classes corresponding to an apparent intermediate mLeptin:mLEP-R complex were selected and used for ab initio reconstruction followed by homogeneous and non-uniform refinement in cryoSPARC, which resulted in an anisotropic cryo-EM map for an 1:2 mLeptin:mLEP-R complex (FSC_0.143_ = 4.43Å with 28,296 particles). An atomic model for a 1:2 mLeptin:LEP-R_IgCRH2FnIII_ complex was created based on the AlphaFold ^45, 46^ prediction for mLEP-R_ECD_ (AF-P48356-F1-model_v1) and the determined mLeptin:mLEP-R_IgCRH2_ and mLEP-R_FnIII_ module crystal structures and fitted in the cryo-EM map via Chimera. The atomic model was further refined via real space refinement in Phenix against the sharpened cryo-EM map using rigid body refinement and coordinate refinement with reference restraints to the starting model and hydrogen-bonding restraints across the site II and site III mLeptin:mLEP-R interface regions. The COSMIC2 webserver ^97^ was used to align projections of the trimeric mLeptin:mLEP-R_IgCRH2_ crystal structure, the 1:2 mLeptin:LEP-R_IgCRH2FnIII_ cryo-EM model, and projections of a 1:1 mLeptin:mLEP-R_ECD_ model based on the AF2-model for mLEP-R_ECD_ and the mLeptin:mLEP-R_CRH2_ crystal structure with high-resolution 2D-classes obtained for the mLeptin:mLEP-R_ECD_ complex sample.

#### mLeptin:mLEP-R_ECD-tGCN4_ complex

The mouse LEP-R ectodomain (mLEP-R_ECD_, residues 1 to 839) fused to a C-terminal trimeric GCN4 (tGNC4) zipper tag via a 5XGGS linker region was secreted from suspension HEK293 FreeStyle cells (ThermoFisher) in the presence of 20 μM kifunensine (Dextra) via PEI-mediated transient transfection as described above. N-terminally His-tagged, refolded mouse Leptin was added to the clarified conditioned medium and the mLeptin:mLEP-R_ECD-tGCN4_ complex was purified via IMAC and a Superose 6 Increase SEC-column with HBS as running buffer. Top fractions were aliquoted and stored at -80 °C until further use. 4 μl of mLeptin:mLEP-R_ECD-tGCN4_ complex at 0.3 mg/mL was applied to a glow-discharged C-Flat 1.2/1.3 300 mesh copper grid (Protochips), blotted for 4.5 s under 99% humidity at 22 °C and plunged into liquid ethane using a GP2 Leica grid plunger. Grids were screened using an JEOL 1400 Plus miscroscope (BECM, Brussels). Data collection was performed using a Titan Krios microscope equipped with a 6k x 4k GATAN K3 BIOQANTUM (eBIC, Didcot) resulting in 13,544 movies with a raw pixel size of 0.829 Å.

Movies were processed via patch-based motion correction and CTF estimation as implemented in cryoSPARC v3.3.1. Particles were extracted with a box size of 448 pixels with 2x binning. Initial 2D classes were obtained by the blob picker function in cryoSPARC, followed by particle picking via TOPAZ as implemented in cryoSPARC. Junk particles were removed by multiple rounds of 2D classification and ab initio 3D classification resulting in a particle set of 141,157 particles. Ab initio 3D classification followed by homogeneous and non-uniform refinement resulted in two cryo-EM maps representing the 3:3 mLeptin:mLEP-R_ECD_ complex. One map displayed better density for the 3:3 mLeptin:mLEP-R_IgCRH2_ core region of the complex and was used for local refinement (FSC_0.143_ resolution = 4.42 Å with 54,708 particles). In the second map for the 3:3 mLeptin:mLEP-R_ECD_ the membrane-proximal regions of the receptor legs are better defined (FSC_0.143_ = 4.60 Å with 37,530 particles). An atomic starting model for the 3:3 mLeptin:LEP-R_ECD_ complex was created based on the AlphaFold prediction for mLEP-R_ECD_ and the determined 3:3 mLeptin:mLEP-R_IgCRH2_ and mLEP-R_FnIII_ module crystal structures and fitted in the cryo-EM map via Chimera. The atomic model was further refined via real space refinement in Phenix against the sharpened cryo-EM map using rigid body refinement and coordinate refinement with reference restraints to the starting model and hydrogen-bonding restraints across the site II and site III mLeptin:mLEPR interface regions. In the cryo-EM maps for the 3:3 mLeptin:LEP-R_ECD_ complex strong additional density was observed at all three site II LEP-R_CRH2_:Leptin interface regions that we interpreted as a trapped Ni-ion coordinated by histidine residues in the LEP-R_CRH2_ module and the N-terminal His-tag present on mLeptin (as shown in **Extended Data Fig. 9a**).

To address the breakage of symmetry in the ring-like core region of the complex and possibly revolve the mLEP-R_CRH2_:mLeptin:mLEP-R_CRH1-IgCRH2_ subcomplex at higher resolution, the pseudo-C3 symmetric volume obtained via local refinement of the 3:3 mLeptin:mLEP-R_tGCN4_ core region (FSC_0.143_ resolution = 4.42 Å with 54,708 particles) was aligned to the pseudo-C3 symmetry axis via the Volume Alignment Tool job in cryoSPARC. The associated particle set was re-extracted without binning (0.829 Å per pixel) and symmetry expanded around the C3 axis resulting in 163,914 particles. Using the molmap function in Chimera, a volume blurred) to 25 Å around one mLEP-R_CRH2_:mLeptin:mLEP-R_!"#$%&!"#’_ subcomplex was generated, and transformed into a mask with the Volume Tools job in cryoSPARC (map threshold = 0.04, dilation radius = 5 Å and soft padding width = 15 Å). Local refinement was performed by limiting the rotation and shift search extent around the original consensus refinement poses to 5° and 5 Å, respectively, in combination with a gaussian prior over the pose/shift magnitudes (with SD_ROT_ = 15 Å and SD_SHIFT_ = 7 Å). The center of mass of the mask was used as a fulcrum point. This approach resulted in a cryo-EM map with an FSC_0.143_ resolution of 4.02 Å in which the crystallographic model for the mLEP-R_CRH2_:mLeptin:mLEP-R) complex was fitted using Chimera and real-space refined in Phenix using reference restraints to the starting model. To enhance high-resolution features the cryo-EM map was sharpened with DeepEMhancer^98^.

#### hLeptin:hLEP-R_ECD-tGCN4_ complex

The human LEP-R ectodomain (hLEP-R_ECD_, residues 1 to 839) fused to a C-terminal trimeric GCN4 (tGNC4) zipper tag via a 5XGGS linker region was co-transfected with N-terminally His-tagged human Leptin in suspension HEK293 FreeStyle cells (ThermoFisher) in the presence of 20 μM kifunensine (Dextra) via PEI-mediated transient transfection. The resulting hLeptin:hLEP-R_ECD-tGCN4_ complex was purified via IMAC and a Superose 6 Increase SEC-column with HBS as running buffer. Top fractions were aliquoted and stored at -80 °C until further use. 4 μl of hLeptin:hLEP-R_ECD-tGCN4_ complex at 0.3 mg.ml^-1^ was applied to a glow-discharged Quantifoil R 0.6/1 300 mesh golden grid coated with graphene (PUXANO), blotted for 1 s (blot force = 1) under 100% humidity at 22 °C and plunged into liquid ethane using an FEI Vitribot Mark IV. Grids were screened using an JEOL 1400 Plus microscope (BECM, Brussels). Data collection was performed using a JEOL cryoARM 300 microscope equipped with a 6k X 4k GATAN K3 detector resulting in 13,755 movies with a raw pixel size of 0.76 Å (BECM, Brussels). Movies were processed via patch-based motion correction and CTF estimation as implemented in cryoSPARC v3.1.0. Particles were extracted with a box size of 560 pixels with 2x binning. Initial 2D-classes were obtained via template-based particle picking using 2D-classes representing top views for a mLeptin:mLEP-RΔ_FnIII-tGCN4_ complex lowpass filtered to 20 Å. The mLEP-RΔ_FnIII-tGCN4_ construct corresponds to a tGCN4-trimerized version of mLEP-R_ECD_ without the membrane-proximal FNIII-module. The cryo-EM dataset for the mLeptin:mLEP-RΔ_FnIII-tGCN4_ complex we collected suffered from severe preferred particle orientations limiting further analysis. The obtained 2D classes for the hLeptin:hLEP-R_ECD-tGCN4_ complex were then used to seed template-based picking and neural network-based particle picking via Topaz 0.2.4. Junk particles were removed by multiple rounds of 2D classification resulting in a particle set of 206,218 particles. High-resolution 2D classes were selected for ab initio 3D classification followed by heterogeneous refinement and non-uniform refinement which resulted in a cryo-EM map that showed well-defined density for a 2:2 hLeptin:hLEP-R_ECD_ subcomplex (91,899 particles, FSC_0.143_ = 5.62 Å). This particle set was further subclassified via ab initio 3D classification and heterogeneous refinement followed by non-uniform refinement which resulted in cryo-EM maps representing closed (29,899 particles, FSC_0.143_ = 6.45 Å) and open (30,353 particles, FSC_0.143_ = 6.84 Å) states of the 3:3 hLeptin:hLEP-R_ECD_ complex.

Starting models model for the 2:2 and 3:3 hLeptin:hLEP-R_ECD_ complexes were created based on the AlphaFold predictions for hLEP-R_ECD_ (AF-P48357-F1-model_v2) and hLeptin (AF-P41159-F1-model_v2), and the determined crystal structures for the mouse 3:3 Leptin:LEP-R_IgCRH2_ complex and human Leptin:LEP-R_CRH2_ complex and fitted in the cryo-EM maps via Chimera. Atomic models were further refined via real space refinement in Phenix against the sharpened cryo-EM maps using rigid body refinement and coordinate refinement with reference restraints to the starting model.

Cryo-EM data collection, processing and refinement statistics are also summarized in **Extended Data Table 3**. All maps are reconstructed in C1 symmetry. Potential duplicate particles were removed within a distance of 150 Å. Reported resolutions are based on the gold-standard Fourier shell correlation (FSC) of 0.143 criterion ^99^ and Fourier shell correlation curves were corrected for the effects of soft masking by high-resolution noise substitution ^100^. Map to model correlations were calculated using phenix.mtriage using the independent half maps as input. Local resolution maps were computed using the blocres algorithm^101^ as implemented in cryoSPARC with an FSC threshold of 0.5. All representations of cryo-EM density and structural models were prepared with ChimeraX ^102^ and PyMol ^103^.

### In silico structure prediction

Structural models for 1:1 mLeptin:mLEP-R_Ig_ and 1:1 mLeptin:mLEP-R_CRH2_ complexes, and for hexameric Leptin:LEP-R_IgCRH2_ assemblies were predicted using the latest version 2.2 of AlphaFold-Multimer implemented through ColabFold ^57, 58^. AlphaFold2-ptm was used for the prediction of mLEP-R_IgCRH2._ Predicted structural models were aligned to the 3:3 mLeptin:mLEP-R_IgCRH2_ crystal structure in Pymol.

### Determination of disulfide bridges by mass spectrometry

Sample preparation: To 50 µL of mLEP-R_ECD_, 5 µL of 1 M Tris-HCl pH 8.0 was added, followed by 5 µL of PNGase F (New England Biolabs, P0704L). The sample was incubated at 37°C for 3h to allow the de-N-glycosylation to proceed. Next, the sample was denatured by adding guanidine hydrochloride to a final concentration of 3M, and stored at room temperature for 20 min. A buffer exchange was performed using a ZebaTM spin desalting column (ThermoFisher Scientific, 89882), with 100 mM Tris-HCl pH 7.5 as destination buffer. Trypsin (Promega, V5117) was added at a 1:25 (w:w) protease/protein ratio, and the sample was incubated overnight at 37°C.

LC-MS: The tryptic digest was analyzed on LC-MS using a C18 column and with H2O, acetonitrile and formic acid as mobile phase constituents. The analysis was performed on a 1290 Infinity UHPLC system coupled to a 6540 Q-TOF mass spectrometer (both Agilent Technologies). The MS instrument was equipped with a AJS-ESI source and operated in positive electrospray ionization mode. The TOF was calibrated prior to the analysis and subsequently operated without utilizing reference masses. Data was collected in centroid mode at a rate of 3 spectra per s in the extended dynamic range mode (2 GHz). MS/MS experiments were performed using data-dependent acquisition (DDA). One survey MS measurement was complemented with 5 data-dependent MS/MS measurements. Doubly, triply and quadruply charged precursor ions were selected based on abundance. After being fragmented twice, a particular m/z value was excluded for 30 s. The quadrupole was operated at medium resolution and the collision energy varied based on m/z. Data acquisition was performed with MassHunter Data Acquisition software, while data processing was performed in MassHunter Qualitative Analysis with BioConfirm add-on (all from Agilent Technologies).

## EXTENDED DATA

**Extended Data Figure 1.**
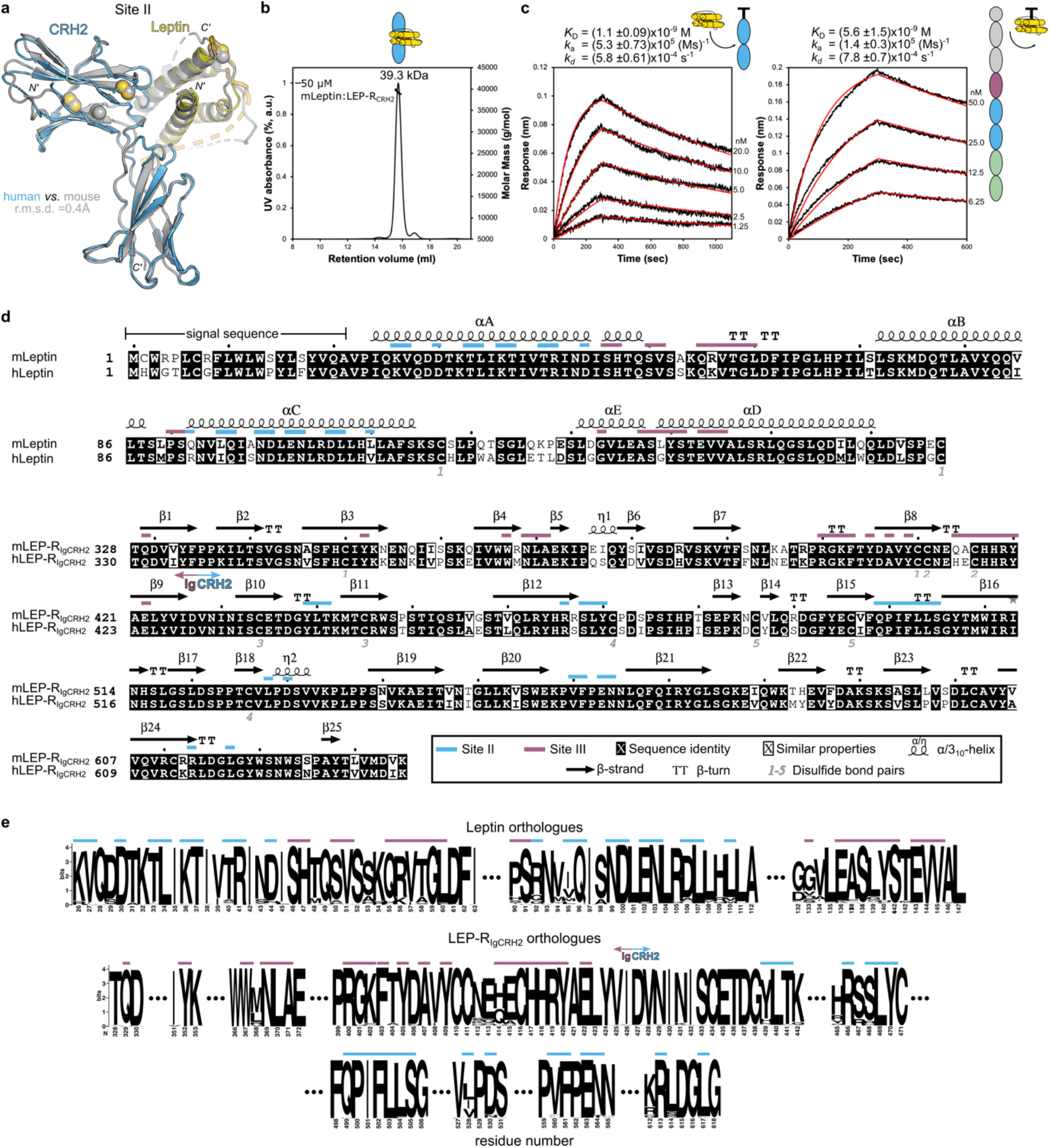
Structural and biophysical characterization of the minimal Leptin:LEP-R recognition complex. **a,** Structural superposition of the glycan-trimmed hLeptin:hLEP-R_CRH2_ complex (CRH2 fragment 428-635 N516Q/C604S in blue; Leptin in yellow) with the mouse complex (colored in gray). The structure of the individual components and the interaction interface thereof is highly similar between the two homologues, consistent with their high sequence identity (83.2% for Leptin, 86.9% for LEP-R_CRH2_). (r.m.s.d.: root mean square deviation of alignment, 161 Cα atoms) **b,** Molar mass determination of the glycan-trimmed mLeptin:mLEP-R_CRH2_: complex using Size Exclusion Chromatography coupled with Multi-Angle Laser Light Scattering (SEC-MALLS). The complex (50 μM) was resolved in a Superdex 200 increase column pre-equilibrated in HBS buffer (20 mM HEPES pH 7.4, 150 mM NaCl). The measured mass (39.3 kDa) approximates the theoretical of 40.3 kDa for 1:1 stoichiometry. **c,** Biolayer interferometry (BLI) sensograms of the mouse Leptin:LEP-R interaction. Left: mLeptin was titrated, as indicated on the side, against the *in vitro* biotinylated mLEP-R_CRH2_ fragment immobilized on Streptavidin sensors *(n = 4)*. Right: mLEP-R_ECD_ was titrated against *in vitro* biotinylated mLeptin *(n = 3)*. Comparison of the kinetic rates and derived affinities suggest that the high-affinity Leptin:LEP-R interaction is attributed to the mLeptin:mLEP-RCRH2 interface, in agreement with previous studies ^37^. **d,** Structurally annotated sequence alignment of mouse with human Leptin (top) and LEP-R_IgCRH2_ fragment (bottom) using ClustalOmega and the ESPript server (http://espript.ibcp.fr/). Site II and III interface residues are highlighted accordingly. Secondary structure elements and disulfide bonds were derived from the crystal structure of the mLeptin:mLEP-R_IgCRH2_ complex determined in this study. **e,** Conservation of the site II and III interface in Leptin and LEP-R orthologues in mammals. Sequence logos were generated using the Weblogo server (http://weblogo.berkeley.edu/logo.cgi) after ClustalOmega multiple sequence alignment of 82 and 73 Leptin and LEP-R orthologues respectively, derived from the Ensembl database (https://www.ensembl.org/index.html). Not fully sequenced orthologues and those not initiating with a Methionine residue were excluded. Interface residues are highlighted as in panel d.

**Extended Data Figure 2.**
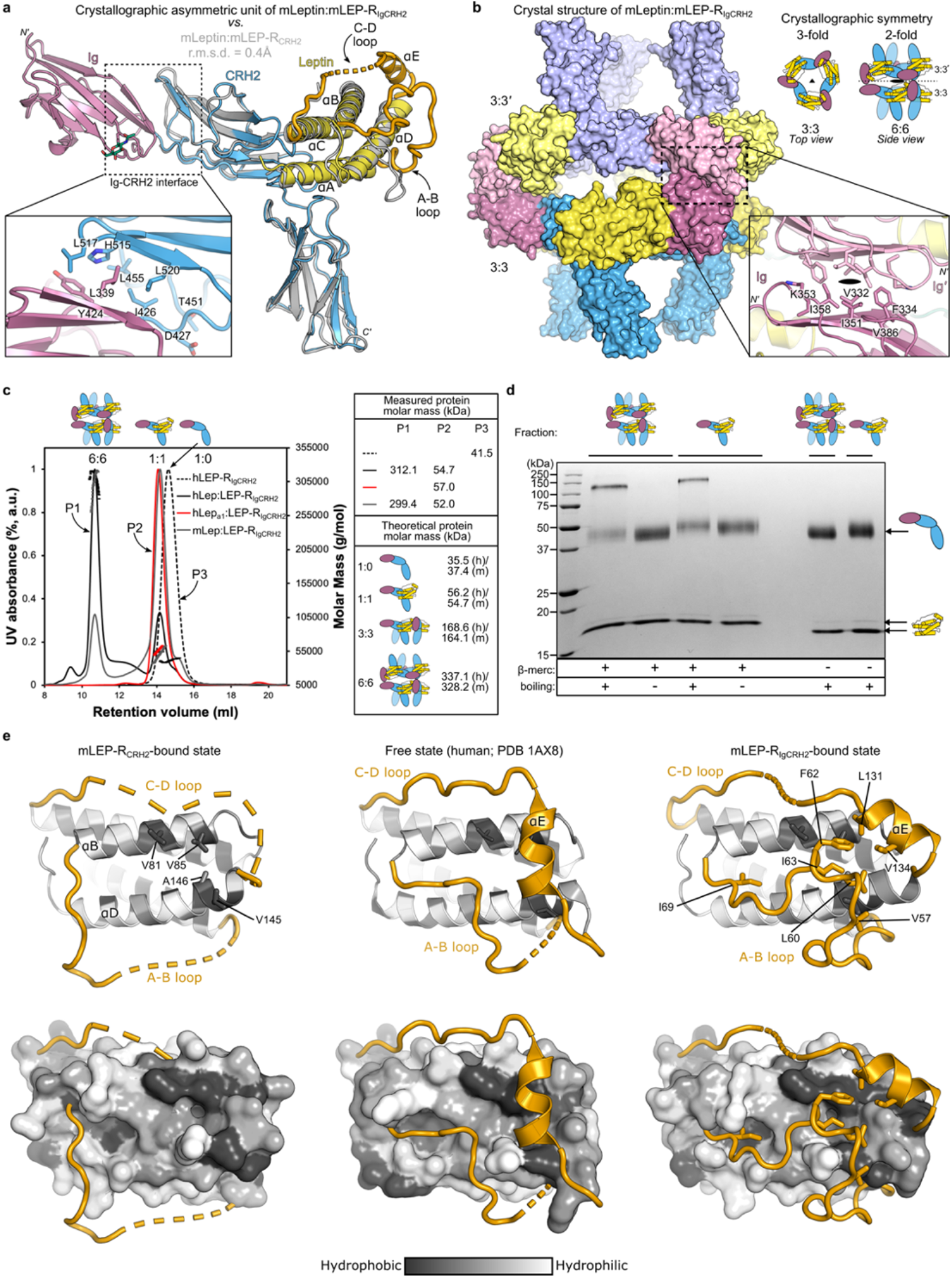
Structural and biophysical characterization of the mLeptin:LEP-R_IgCRH2_ oligomeric assembly. **a,** The asymmetric unit of the glycan-trimmed mLeptin:mLEP-R_IgCRH2_ crystal structure resolved at 2.9Å (coloured as in Fig. 1a) superimposed to the mLeptin:mLEP-R_CRH2_ structure (grey) (r.m.s.d. of alignment =0.4Å, 257 Cα atoms). Interacting residues at the Ig-CRH2 interface are highlighted in the insert. A glycan chain (green sticks) was resolved at position Asn393 of the Ig domain. **b,** The oligomeric assembly of mLeptin:mLEP-R_IgCRH2_ (left) derived from the 3- and 2-fold crystallographic symmetry operations as detailed on the right (space group H 3 2). Symmetry axis are indicated by a triangle and an oval respectively. The assembly comprised of a 3:3 Leptin:LEP-R_IgCRH2_ stoichiometric complex through site II and site III contacts, and an artifactual head-to-head dimerization thereof, resulting in a 6:6 complex. The head-to-head dimer associates purely by mirroring Ig-Ig’ contacts at an exposed hydrophobic patch shown in the insert. The LEP-R N-terminal domain is expected to disrupt this interface (N-termini indicated). **c,** In solution molar mass determination of LEP-R_IgCRH2_ and complexes thereof with Leptin and the Leptin antagonist a1 (^140^ST^141^/AA) ^39, 47^ using SEC-MALLS (Superdex 200 increase column). Protein molar masses are plotted after protein conjugate analysis of the glycosylated complexes. Experimentally determined and theoretical masses are given on the right. h: human; m: mouse complexes. **d,** SDS-PAGE (17%) of isolated 6:6 and 1:1 Leptin:LEP-R_IgCRH2_ complexes under different sample preparation conditions. Multimeric artifacts are observed only with the combination of reduction and boiling, raising awareness for sample preparation for determining stoichiometry from SDS-PAGE. Leptin migrates slower under reducing conditions. (β-merc: β-mercaptoethanol) **e,** Conformational plasticity in the loops of Leptin. The partially resolved A-B and C-D loops of Leptin (orange) are shown relatively to the main four-helix-bundle body coloured according to the Eisenberg hydrophobicity-gradient scale^104^ at the mLEP-RCRH2-bound (left), free ^1^ (middle; pdb: 1ax8; hLeptin) and mLEP-R_Ig_-bound state (right). A hydrophobic hub is formed at the surface of helices αB and αD of the four-helix-bundle by the indicated residues in grey. When not bound to LEP-R_Ig_, the long loops of Leptin are dynamic and may transiently interact with this hub. At the free state, the α-helix E was stabilized on the same hub due to crystal packing. At the LEP-R_Ig_-bound state, the hydrophobic hub is occupied by the A-B loop which in turn stabilizes the α-helix E in a V-configuration. This loop restructuring activates the site III on Leptin.

**Extended Data Figure 3.**
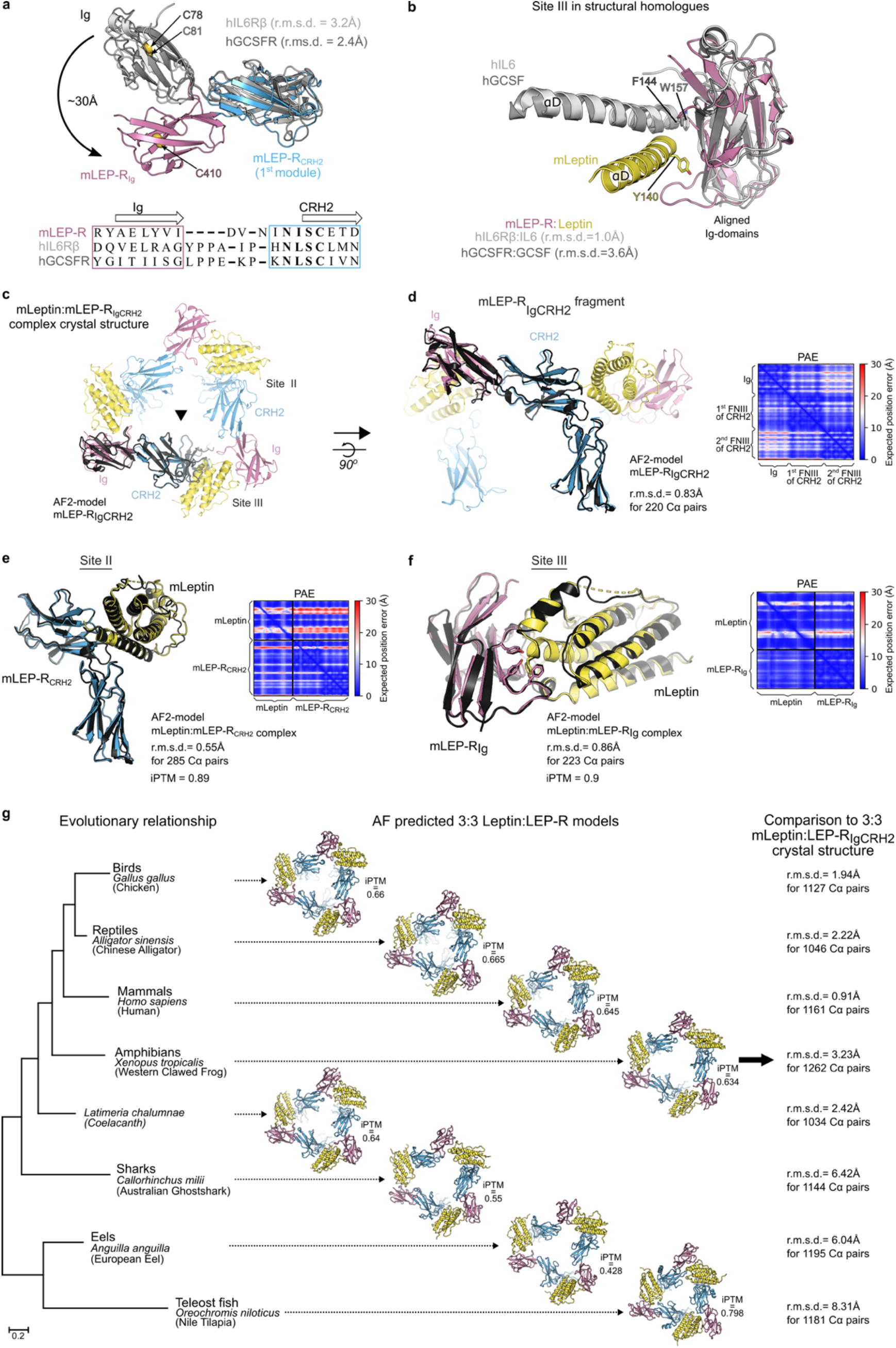
Evolutionary and co-evolutionary features of LEP-R_IgCRH2_ and its interaction interfaces with Leptin. **a,** Structural superposition (top) and sequence alignment (bottom) of the LEP-R_IgCRH2_ domains with those of human gp130 (IL6Rβ) ^27^ and GCSFR ^28^ in their cytokine-bound state (pdb: 1p9m, 2d9q respectively). After alignment at the first FNIII domain of the CRH2 module (r.m.s.d. indicated; 75 and 60 Cα atoms respectively), the shift distance was determined between the highly conserved cysteine residues of the Ig motif, as indicated. Structure-based sequence alignment was performed using UCSF Chimera and shows a shorter linker between the Ig and CRH2 domains for LEP-R relatively to its structural homologues. **b,** Structural superposition of the site III interface in evolutionary relatives. For visualization purposes only the α-helix D is shown for mLeptin, hIL6 and hGCSF, featuring the aromatic blueprint of site III in sticks. IL6Rβ:IL6 and GCSFR:GCSF were aligned to the Ig domain of mLeptin:LEP-R_IgCRH2_ (r.m.s.d. indicated; 41 and 59 Cα atoms respectively). **c-f,** Structural superposition of the crystallographically-distilled mLeptin:mLEP-R_IgCRH2_ assembly with AlphaFold models, as indicated, accompanied with predicted Aligned Error plots (PAE) of each prediction. AlphaFold2-ptm was used for the prediction of mLEP-R_IgCRH2_ (panels c-d) and AlphaFold-Multimer (version 2.2) for the prediction of the mLeptin:mLEP-R_IgCRH2_ interaction interfaces (panels e-f). **g,** Evolutionary conservation of the Leptin:LEP-R_IgCRH2_ hexameric assembly across the animal phylogenetic spectrum, tested by AlphaFold-Multimer (version 2.2, with associated ipTM scores) ^57, 58^ with orthologous 3:3 pairings of Leptin and LEP-R sequences harvested from UniProt. Drawing the branching of jawed vertebrates from the coelacanth phylogenetic tree ^105^, birds are represented by the chicken *G. gallus* Leptin:LEP-R complex (respectively UniProt codes O42164 and Q9I8V6), reptiles by the alligator *A. sinensis* (A0A1U8DFZ0 and A0A3Q0H8E2), mammals by human *H. sapiens* (P41159 and P48357), amphibians by *X. tropicalis* (A0A803KEF7 and F6RVW6), lobe-finned fishes by the living fossil *Coelacanth L. chalumnae* (H3AP27 and H3AG22), cartilaginous fishes by the Australian ghostshark *C. milii* (A0A4W3H871 and V9KNM1), and ray-finned fishes by the eel *A. anguilla* (A0A0C7AV37 and A0A0C7AV33) and tilapia *O. niloticus* (I3KCE8 and A0A067Z8Z1). Sequence evolutionary relationships were based on LEP_R orthologues and were adapted from Londraville et al., 2017 ^106^. Complexes were visualized and superposed to the X-ray structure of mLeptin:mLEP-R_IgCRH2_ by PyMOL 2.3.4 (www.pymol.org) with listed r.m.s.d fit to the 3:3 complex.

**Extended Data Figure 4.**
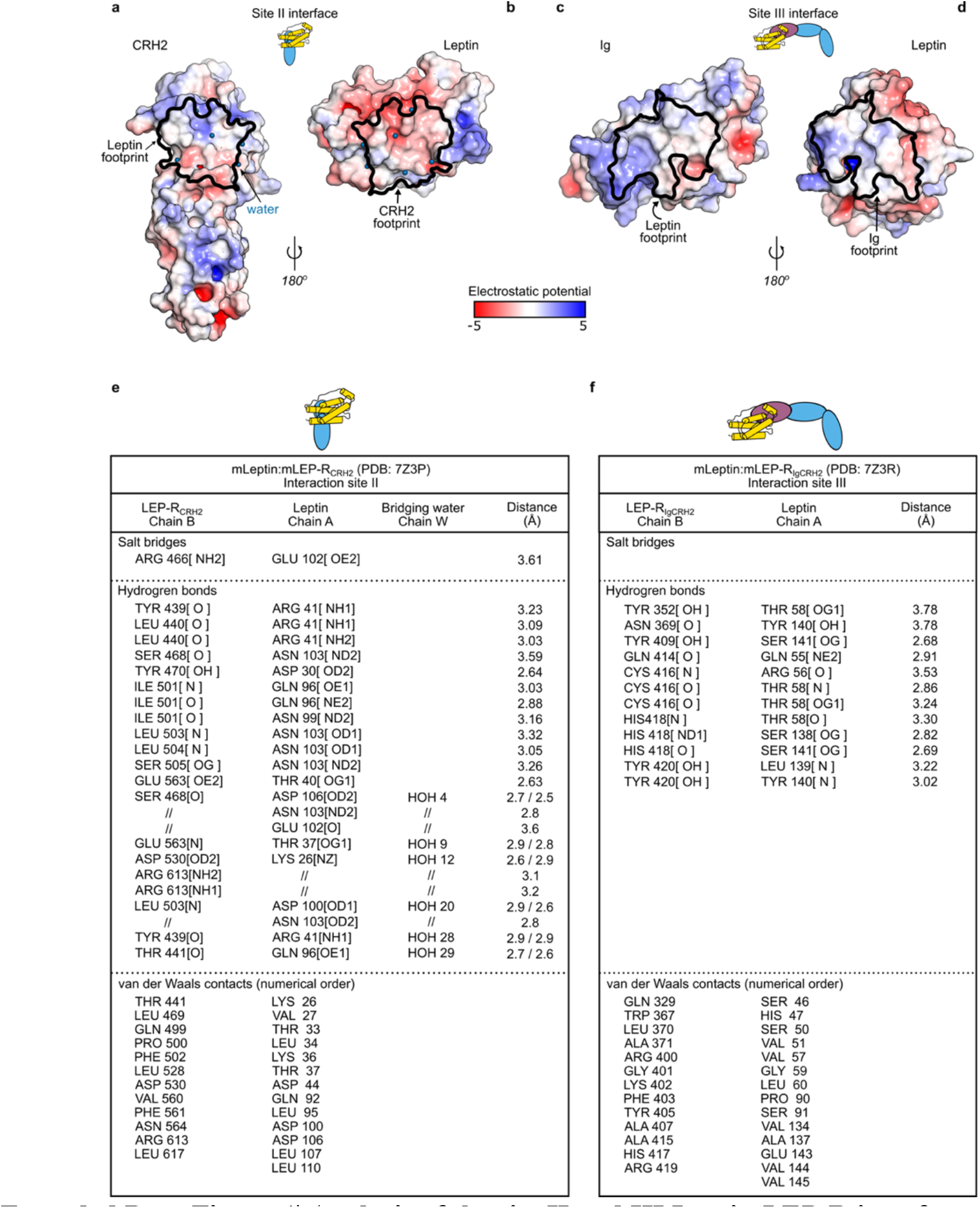
Analysis of the site II and III Leptin:LEP-R interfaces. **a-b**, Site II interface footprint on LEP-R_CRH2_ (a) and Leptin (b) mapped on an electrostatic potential-gradient surface representation of the individual components of the mLeptin:mLEP-R_CRH2_ complex. **c-d,** Site III interface footprint on LEP-R_Ig_ (c) and Leptin (d), as in a-b, in the mLeptin:mLEP-R_IgCRH2_ assembly. **e-f,** Site II (e) and Site III (f) interacting residues in the mLeptin:mLEP-R(Ig)CRH2 complexes as determined by the PISA server (https://www.ebi.ac.uk/msd-srv/prot_int/pistart.html) and manual validation.

**Extended Data Figure 5.**
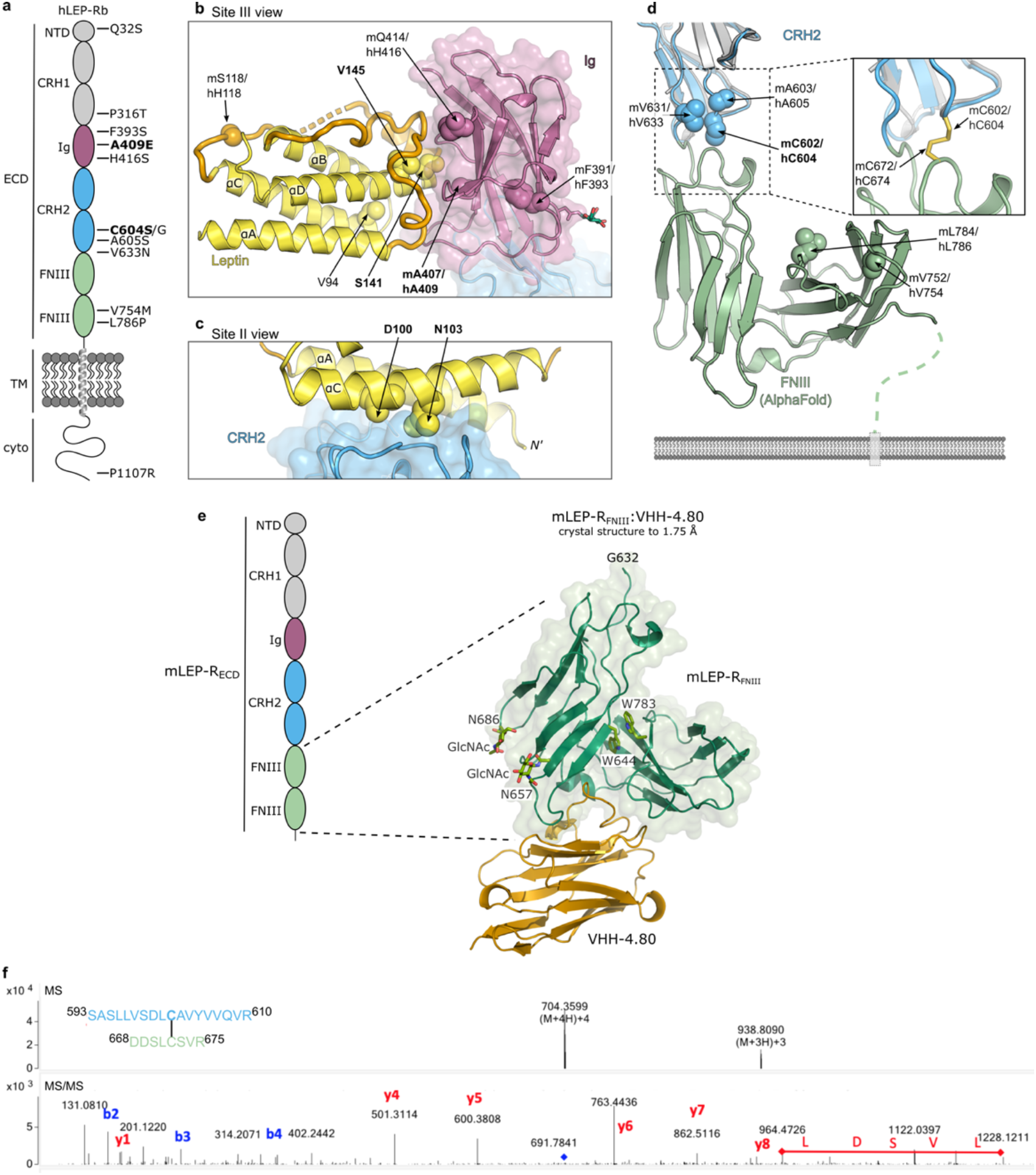
Map of naturally occurring mutations likely related to obesity on the Leptin:receptor assembly. **a,** Domain distribution of missense variants identified in the human LEP-R gene of obese individuals (**Extended Data Table 2**), that have no currently known effect on protein secretion and stability. Functionally validated pathogenic mutations are highlighted in bold. b-c, Localization of mutations in the LEP and LEP-R genes (**Extended Data Table 2**) in the assembly. Mutated residues are shown as spheres. Functionally validated pathogenic mutations are highlighted in bold. d, Structural superposition of the crystallographically-resolved hLEP-R_CRH2_ domain (gray) with the AlphaFold model of hLEP-R (r.m.s.d.=0.669Å, 171 Cα atoms). Residues found in variants likely related to obesity are annotated. A disulfide bridge is predicted to be formed between the CRH2 and FNIII domains (Cys604-Cys674), as shown in the insert. Both residues are functionally important ^22, 25^. The disulfide bridge was confirmed by peptide mapping for mLEP-R_ECD_, as shown in panel f. Cysteine 602 (604 in human) was mutated to serine in FNIII-deletion permutations of LEP-R in this study to prevent artificial disulfide-linked clusters. e, Cartoon representation of the determined crystal structure of for mLEP-RFnIII:VHH-4.80 complex. Evolutionary conserved Trp across the interdomain interface and crystallographically observed N-linked glycosylation sites are shown as sticks. f, Mass spectrometric confirmation of the disulfide bridge between residues Cys602-Cys672 of the CRH2 and FNIII domains respectively in mLEP-R_ECD_. The MS (top) and MS/MS spectra (bottom) are shown for the disulfide-linked peptides that are indicated on the left.

**Extended Data Figure 6.**
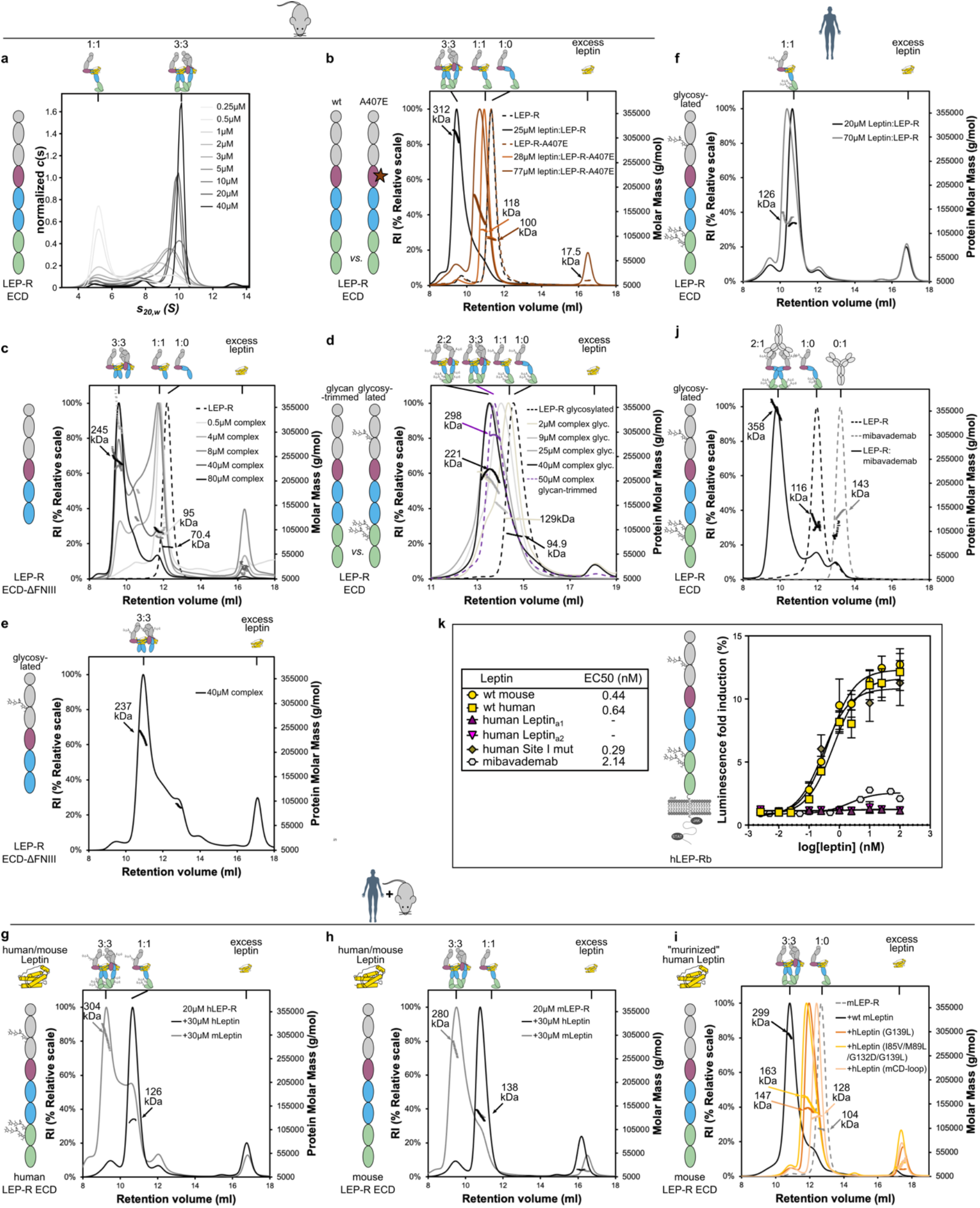
Biophysical and functional characterization of LEP-R complexes. **a-e,** Biophysical analysis of mouse Leptin:LEP-R_ECD_ in solution. **a,** Sedimentation coefficient distributions c(s) for different concentrations of glycan-trimmed mLeptin:mLEP-R_ECD_. **b,** SEC-MALLS analysis (Superdex 200 increase column) of glycan-trimmed wild type mLEP-R_ECD_ and mLEP-R_ECD_-A407 mutant and their complexes with mLeptin. **c and e,** SEC-MALLS analysis (Superdex 200 increase column) of glycan-trimmed (panel c) or glycosylated (panel e) mLEP-R_ECD_-ΔFNIII and their complexes with mLeptin. **d,** SEC-MALLS analysis (Superose 6 increase column) of glycosylated mLEP-R_ECD_ and its complexes with mLeptin, as a function of complex concentration. As a control, the glycan-trimmed complex is given at a single concentration (purple). Formation of mainly oligomeric species is reached at approximately 25μM for both states (see also panel b). **f, j, k,** Characterization of human LEP-R complexes. **f,** SEC-MALLS analysis (Superdex 200 increase column) of glycosylated hLEP-R_ECD_ and its complexes with hLeptin. **j,** SEC-MALLS analysis (Superdex 200 increase column) of glycosylated hLEP-R_ECD_ and its complexes with the agonistic antibody mibavademab^53^. **k,** Activation of hLEP-Rb by Leptin and variants, as well as by the agonistic antibody mibavademab, probed by a STAT3-responsive luciferase reporter in HEK293T cells. *n=3* **g, h, i,** Interspecies cross-reactivity probed with SEC-MALLS (Superdex 200 increase column). **g,** Complexes of hLEP-R_ECD_ with mouse and human Leptin. **h,** Complexes of glycan-trimmed mLEP-R_ECD_ with mouse and human Leptin. **i,** Complexes of glycan-trimmed mLEP-R_ECD_ with “murinized” human Leptin. Specifically, residues close to the Site III interface that differ between mouse and human Leptin (Extended Data Fig. 1d) were mutated on hLeptin to the corresponding of the mouse homologue (i.e. CD-loop residues 118-129; G139L; I85V/M89L/G132D/G139L).

**Extended Data Figure 7.**
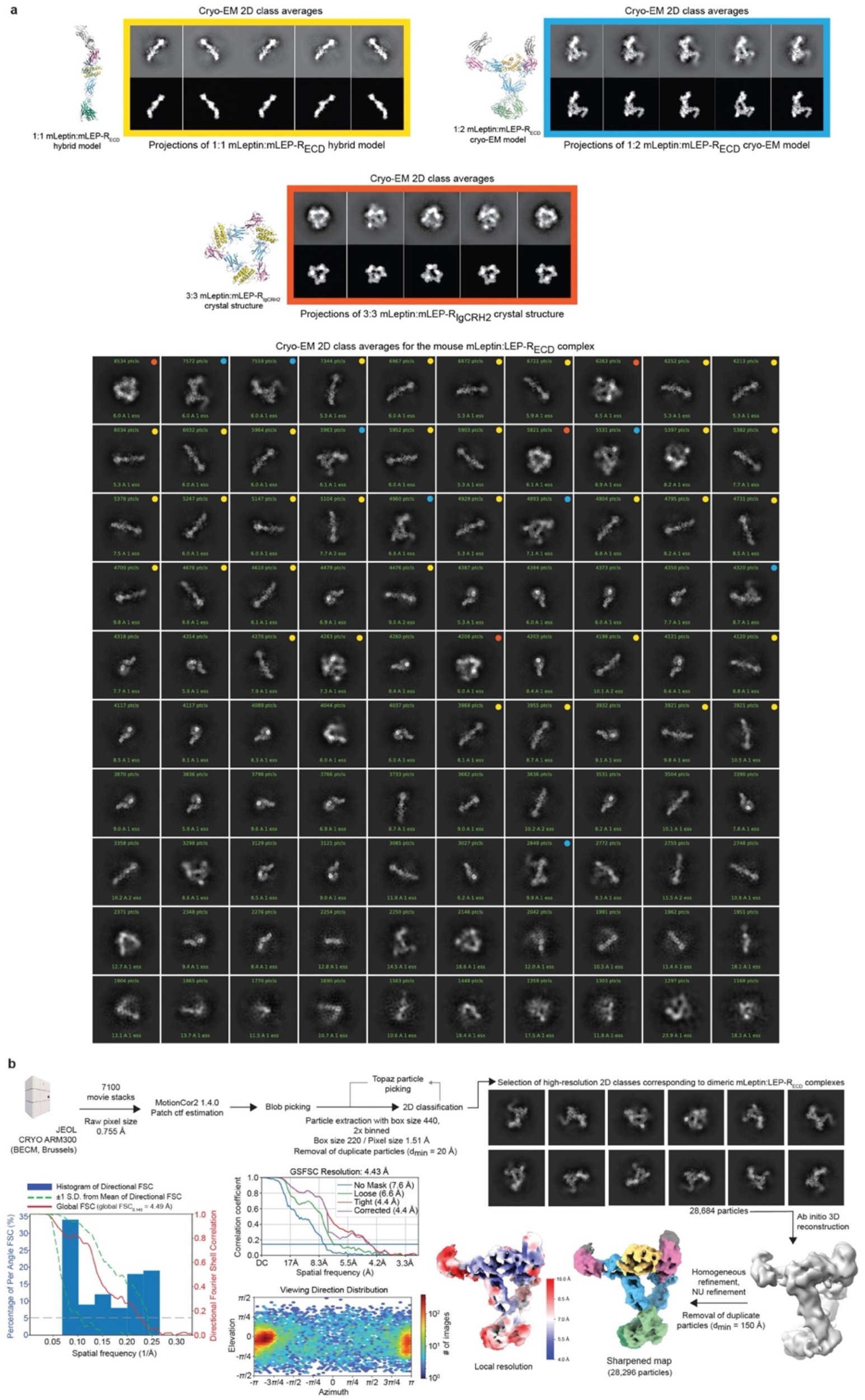
Cryo-EM analysis of mLep:mLEP-R_ECD_ complexes. **a.** The COSMIC^2^ webserver was used to align projections of the trimeric mLeptin:mLEP-R_IgCRH2_ crystal structure, the 1:2 mLeptin:LEP-R_IgCRH2FnIII_ cryo-EM model, and projections of a 1:1 mLeptin:mLEP-R_ECD_ model (based on the Alphafold2-model for mLEP-R_ECD_ and the mLeptin:mLEP-R_CRH2_ crystal structure) with high-resolution 2D-classes obtained for the mLeptin:mLEP-R_ECD_ complex sample. In the gallery 2D cryo-EM classes matching projections are indicated with a colored dot. **b**, Cryo-EM workflow towards the reconstruction of the 1:2 mLeptin:LEP-R_ECD_ complex. We note that 3D reconstruction of the remaining two observed states was hindered due to preferred orientation.

**Extended Data Figure 8.**
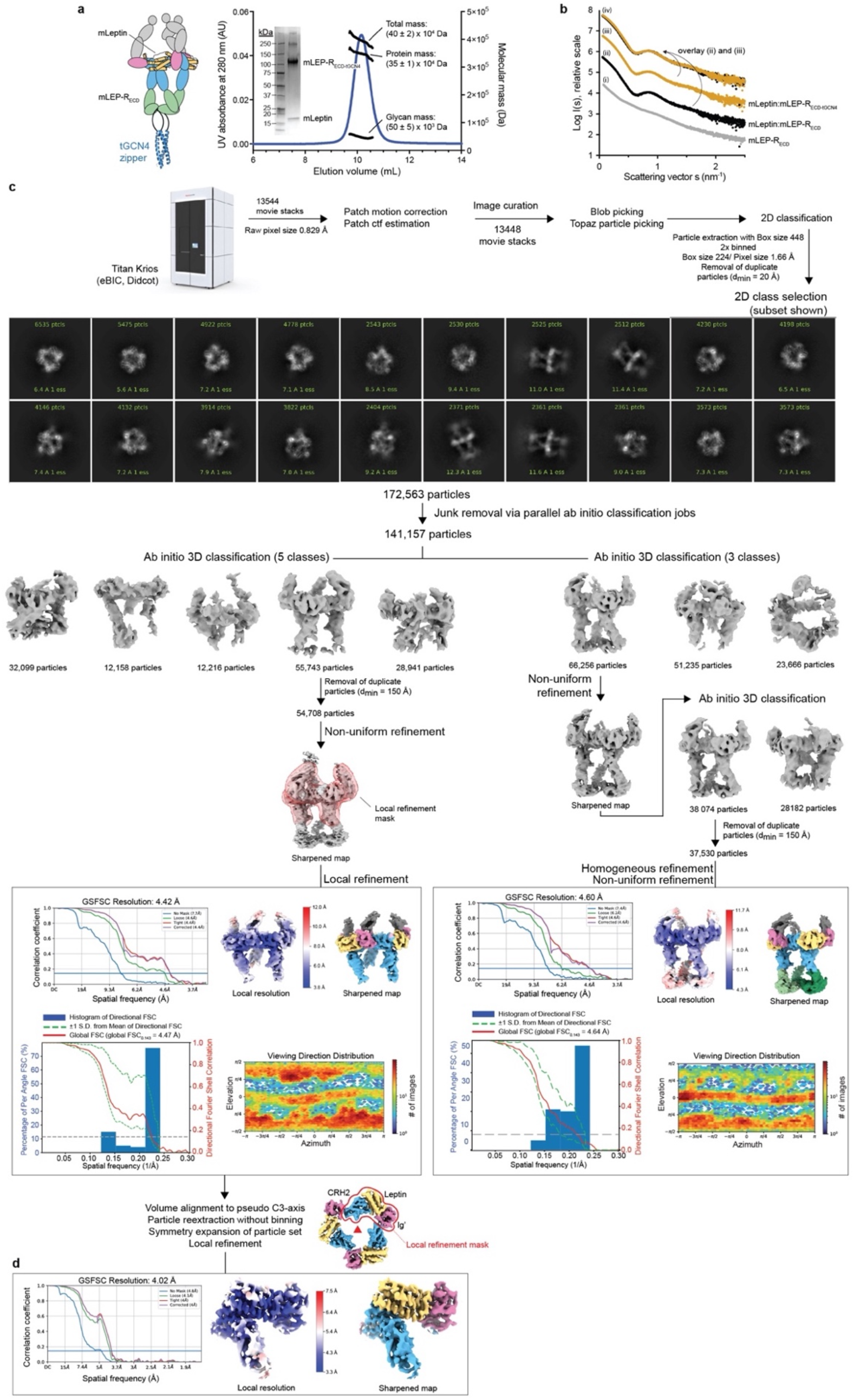
Cryo-EM data analysis workflow for the mLeptin:mLEP-R_tGCN4_ complex. **a,** SEC-MALLS analysis of the glycosylated mLeptin:mLEP-R_ECD-tGCN4_ complex. The SEC-elution profile is plotted as the ultraviolet absorbance at 280 nm (left Y-axis) in function of elution volume. The total, protein and glycan molecular mass (right Y-axis) as determined by MALLS are reported as the number average molecular mass (and s.d.) **b,** SAXS scattering curves of EndoH-treated samples for mLEP-R_ECD_ and mLeptin:mLEP-R_ECD_ complexes plotted as the scattered intensity in function of scattering vector *s* = 4*π*sin*θ*/*λ. (i)* mLEP-R_ECD_ (grey), (*ii*) non-stabilized mLeptin:mLEP-R_ECD_ complex (black), (*iii*) stabilized mLeptin:mLEP-R_ECD-tGCN4_ complex (yellow), *(iv)* scaled and overlayed scattering profiles for mLeptin:mLEP-R_ECD_ and mLeptin:mLEP-R_ECD-tGCN4_. **c.** Cryo-EM data processing workflow in cryoSPARC for the glycosylated mLeptin:mLEP-R_ECD_-ECD-tGCN4 complex. NU: non-uniform refinement.

**Extended Data Figure 9.**
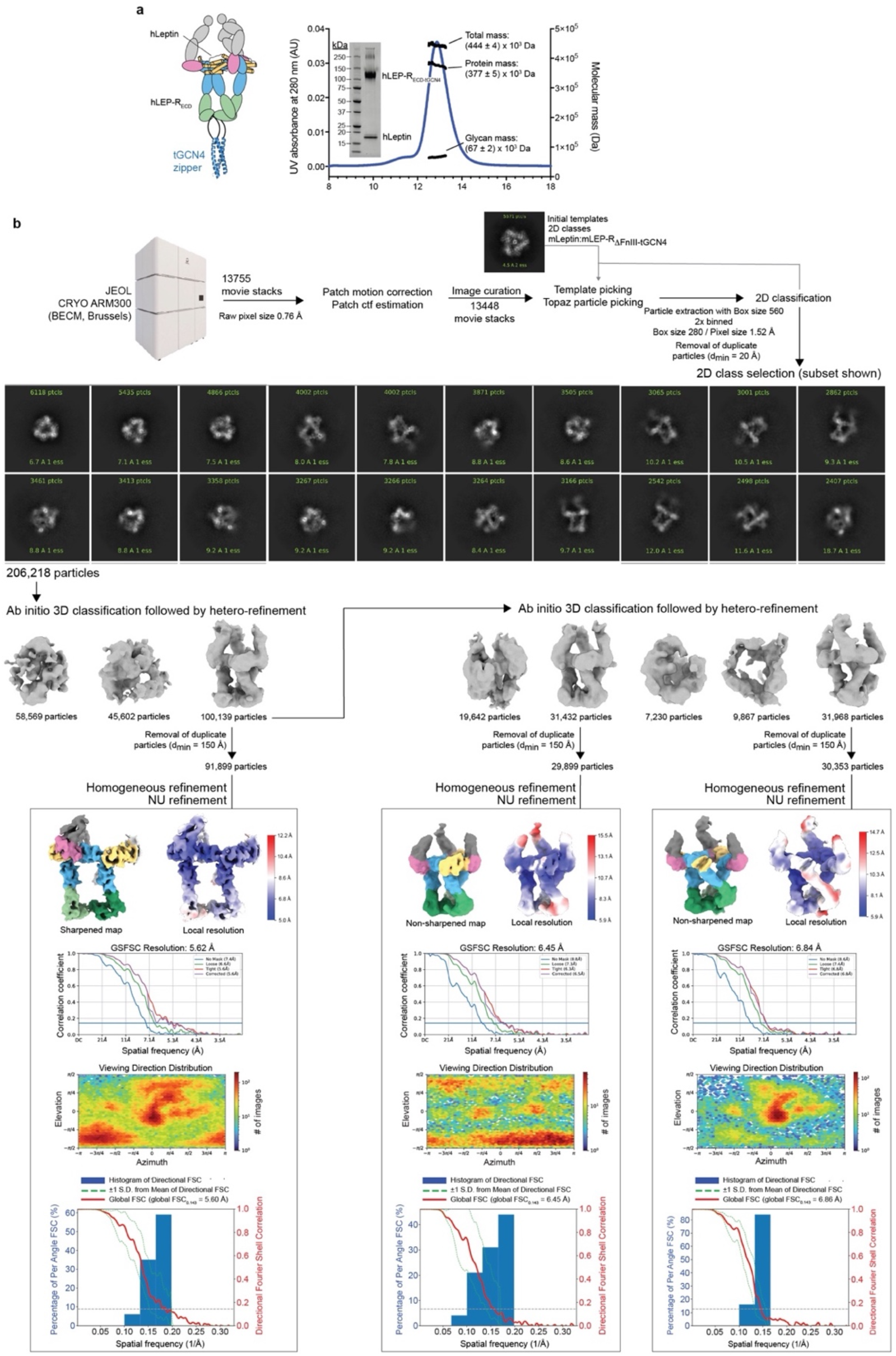
Cryo-EM data analysis workflow for the hLeptin:hLEP-R_tGCN4_ complex. **a,** SEC-MALLS analysis of the glycosylated hLeptin:hLEP-R_ECD-tGCN4_ complex. The SEC-elution profile is plotted as the ultraviolet absorbance at 280 nm (left Y-axis) in function of elution volume. The total, protein and glycan molecular mass (right Y-axis) as determined by MALLS are reported as the number average molecular mass (and s.d.) **b,** Cryo-EM data processing workflow in cryoSPARC for the glycosylated hLeptin:hLEP-R_ECD-tGCN4_ complex. NU: non-uniform refinement.

**Extended Data Figure 10.**
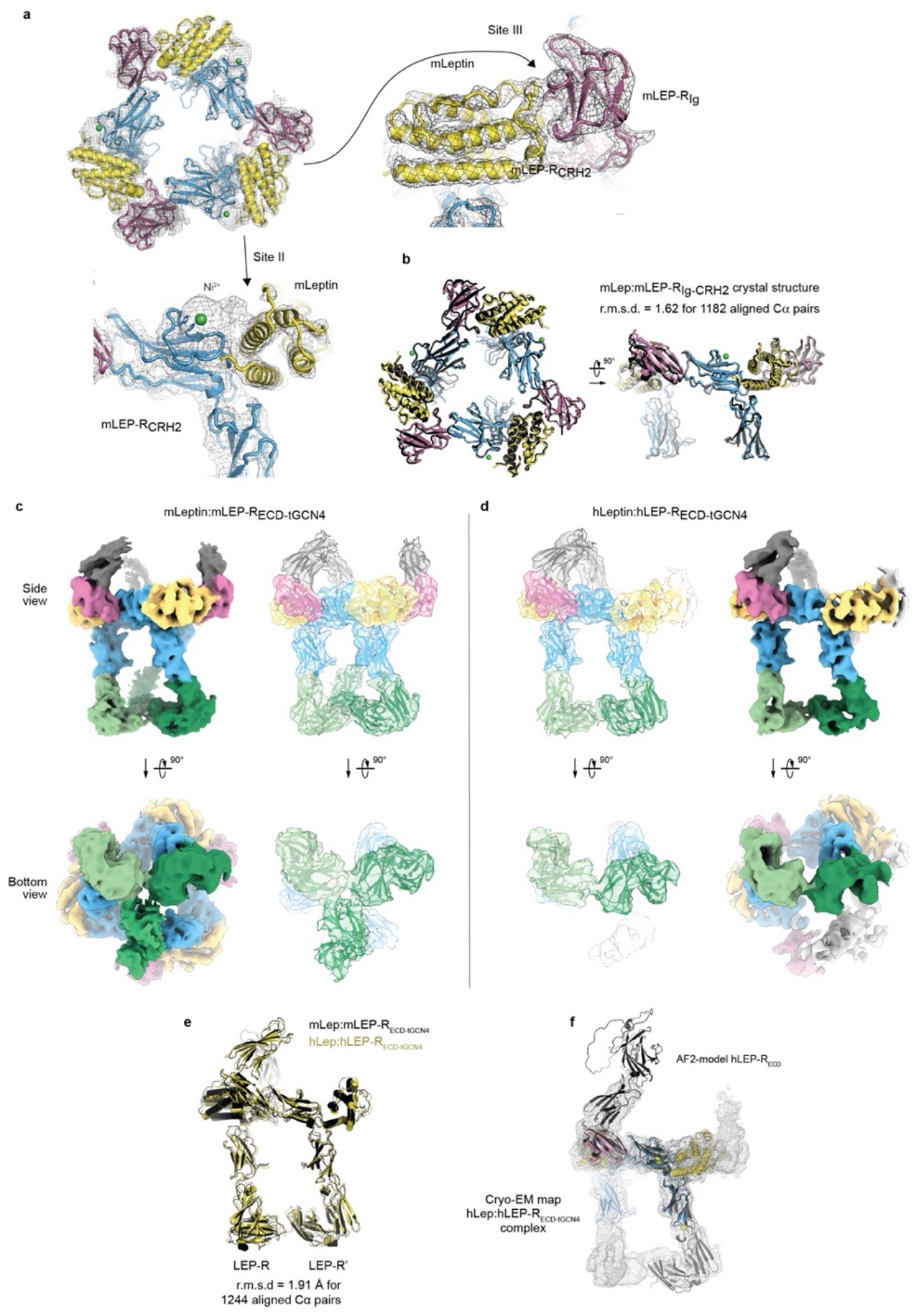
Cryo-EM data analysis. **a,** Real-space refined atomic model for the trimeric mLeptin:mLEP-R_IgCRH2_ core region overlayed with the sharpened cryo-EM map in C1 symmetry for the mLeptin:mLEP-R_ECD-tGCN4_ complex following local map refinement. The cryo-EM map is contoured at 0.313 V. In the cryo-EM map strong additional density is observed at all three LEP-RCRH2:Leptin interface regions (site II) that likely corresponds to a trapped Ni-ion coordinated by histidine residues in the LEP-R_CRH2_ module and the N-terminal His-tag present on mLeptin. **b,** Structural superposition of the crystallographically-distilled 3:3 mLeptin:mLEP-R_IgCRH2_ assembly (black) with the real-space refined mLeptin:mLEP-R_IgCRH2_ assembly via cryo-EM. **c-d,** Cryo-EM maps and fitted atomic models for the mouse and human Leptin:LEP-R_ECD-tGCN4_ complexes. The sharpened cryo-EM maps are colored per zone with Leptin in yellow, the CRH1 module in grey, the Ig domain in magenta, the CRH2 module in blue and the FnIII module in green. The fitted atomic model models are shown as a cartoon overlayed with the cryo-EM maps as a transparent volume. The mouse and human cryo-EM maps are contoured at 0.178 V and 0.550 V, respectively. **e**, Structural superposition of the real-space refined atomic models for the mouse and human Leptin:LEP-R_ECD-tGCN4_ complexes. **f**, Structural superposition of the real-space refined model for the hLeptin:hLEP-R_IgCRH2_ core with the structural prediction for hLEP-R_ECD_ via AlphaFold2 overlayed with the sharpened cryo-EM map for the hLeptin:hLEP-R_ECD-tGCN4_ complex.

**Extended Data Figure 11.**
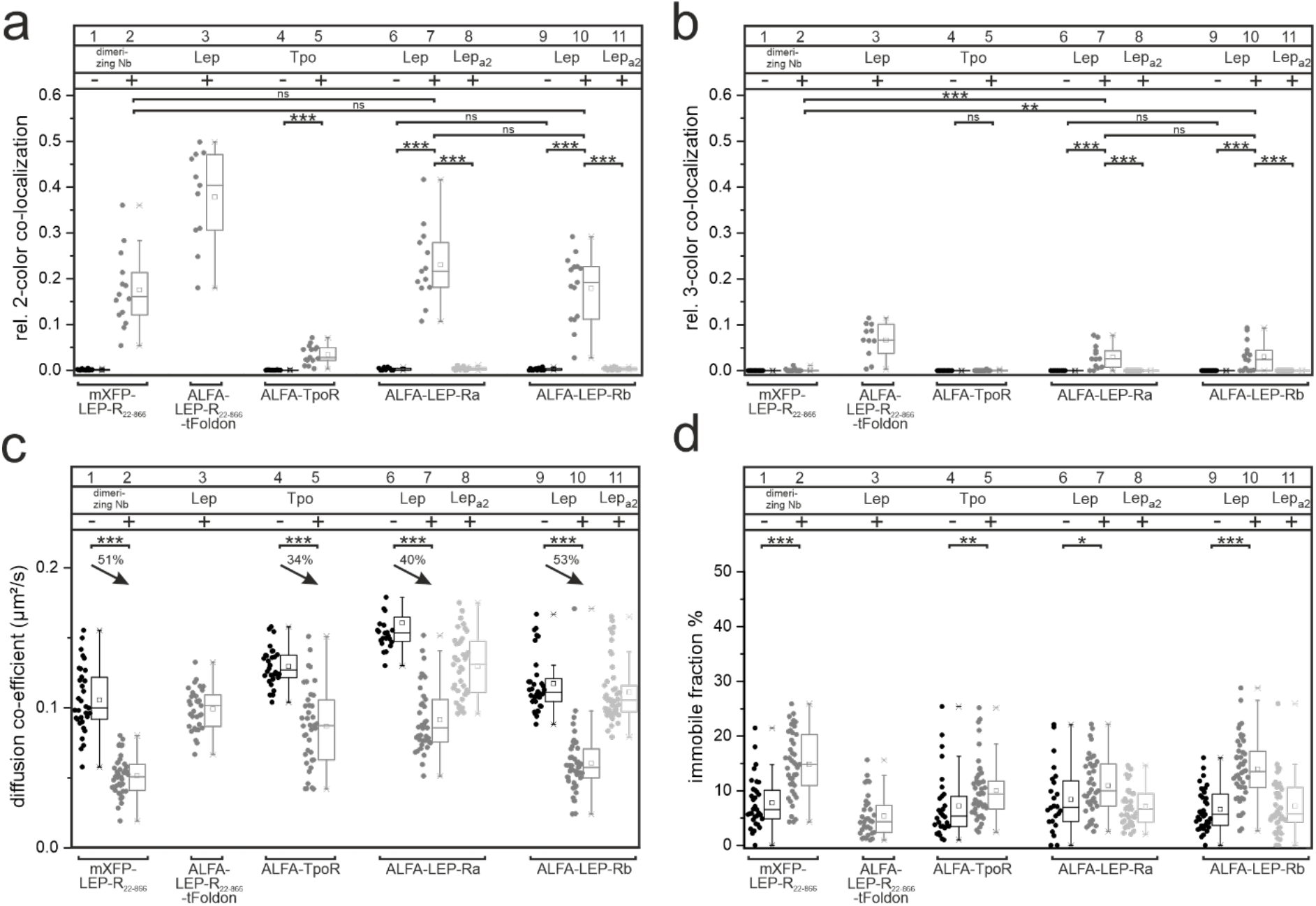
Receptor co-localization at the cell surface probed by smTIRFM. **a,** Dual-color (2C) co-tracking. **b,** Triple-color (3C) co-tracking. **c,** Diffusion properties; arrows indicate percentage of diffusional decrease **d,** Percentage of immobile fraction.

**Extended Data Table 1.**
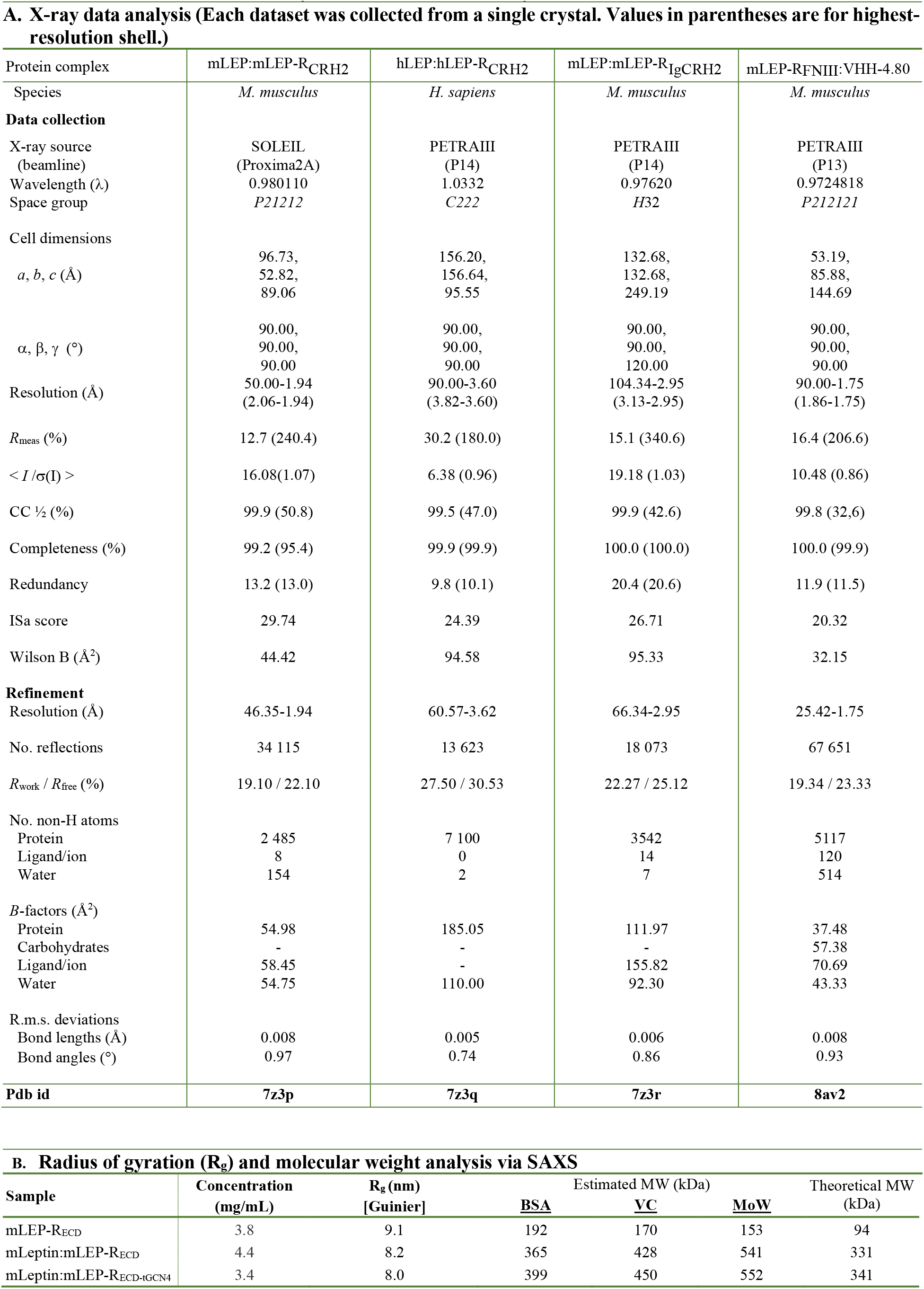
X-ray and SAXS data analysis.

**Extended Data Table 2.**
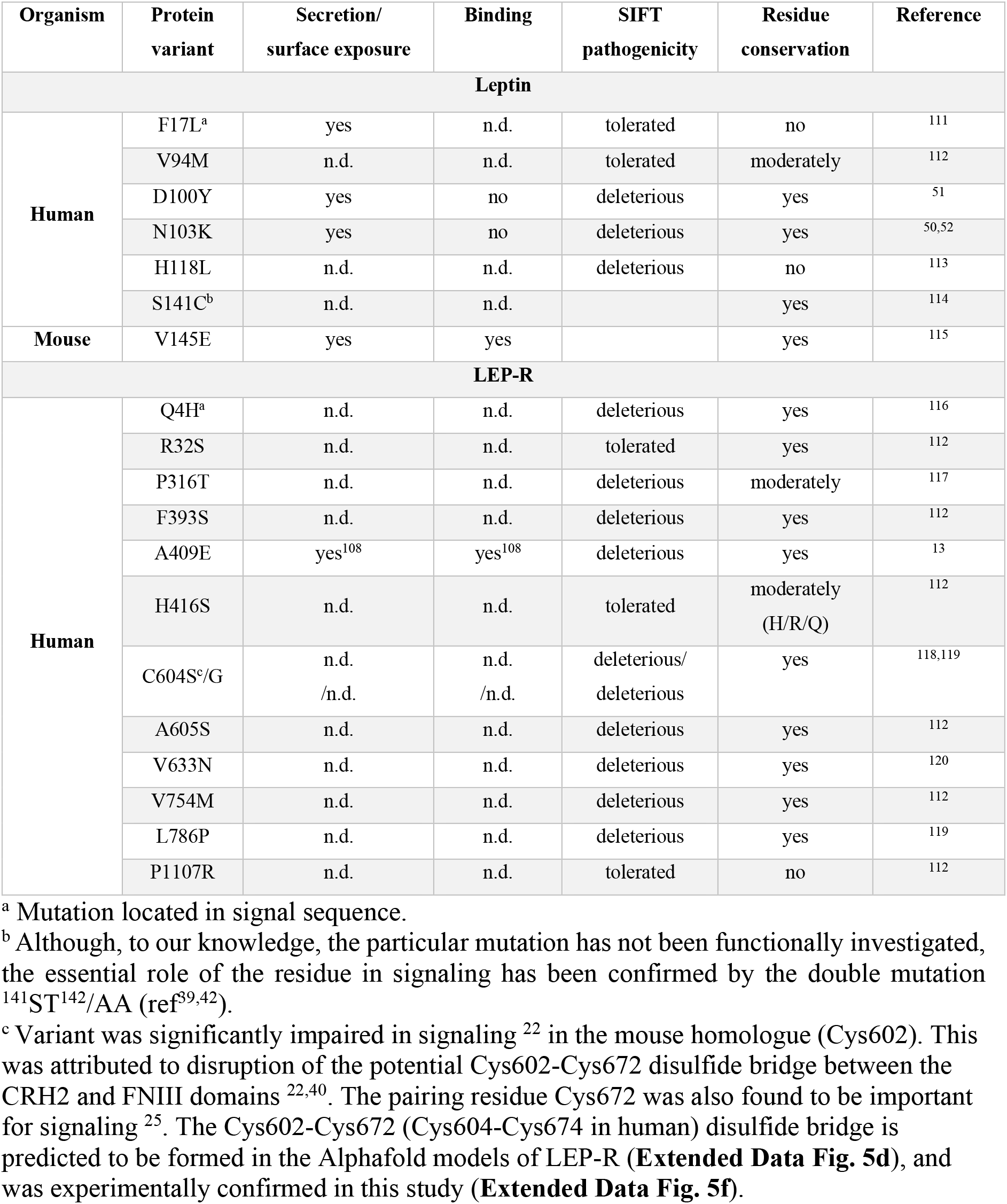
Missense mutations in obese individuals with no apparent/currently known effect in protein secretion and stability. Pathogenic mutations with known impaired expression/secretion/stability ^107–109^ were excluded from this list and were recently detailed in *refs*^12, 49^. As functional validation of most mutations is yet to be provided, the *in silico* prediction of the pathogenicity level using Sorting Intolerant From Tolerant (SIFT) ^110^ program is given.

**Extended Data Table 3.**
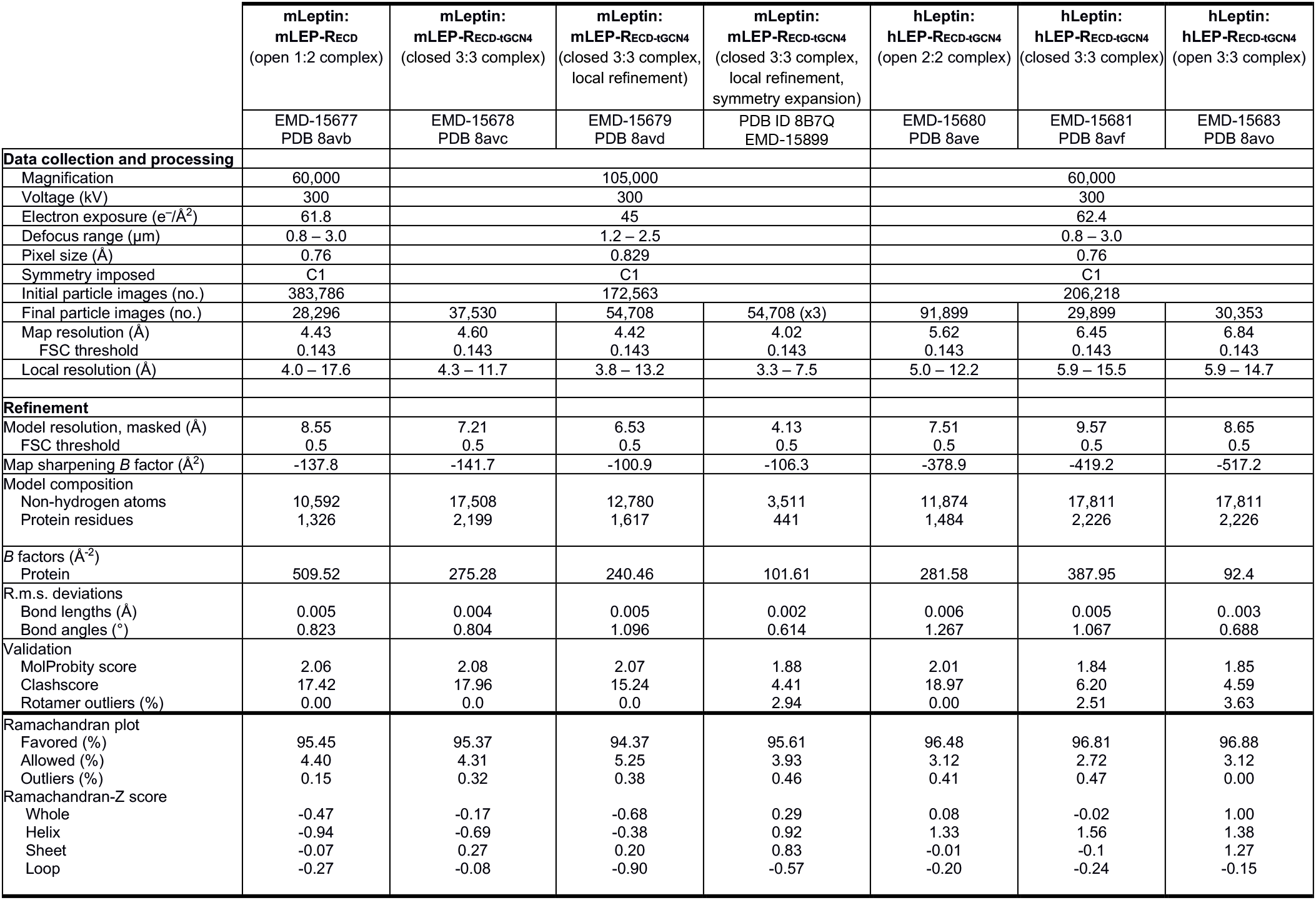
Cryo-EM data collection, processing, refinement and validation statistics.

**Supplementary Movie 1 | mLep:mLEP-Rb assembly in living cells**

Co-localization of mLEP-Rb in the absence (left) and presence of ligand (middle: 2-color co-localization; right: 3-color co-localization, 10 nM mLeptin, 10 min). Red, blue and grey signals correspond to receptors labeled with ^Cy3B^NB, ^Atto643^NB, and ^Dy752^ NB, respectively. Acquisition frame rate: 33 Hz, Playback: real time

## Notes

### Competing Interest Statement

The authors have declared no competing interest.

