## Supplementary Table 1 for "Mechanism of receptor assembly via the pleiotropic adipokine Leptin"

**Supplementary Table 1. List of protein expression constructs used in this study.**

| Protein (segment) | Construct | Expression system | Application |
|---|---|---|---|
| Mouse LEP-$R_{CRH2}$ (426-633; N514Q/C602S) | pCAGGS_ss_His_TEV_mCHR2* (594) | FreeStyle™ 293-F | X-ray crystallography (mLeptin:mLEP-$R_{CRH2}$ assembly), SEC-MALS (Ext. Data Fig. 1b) |
| Mouse Leptin wt (22-167) | pTwist_ss_His_mLEP (586) | FreeStyle™ 293-F | X-ray crystallography (mLeptin:mLEP-$R_{CRH2}$ assembly), SEC-MALS (Ext. Data Fig. 2c) |
| Human LEP-$R_{CRH2}$ (428-635; N516Q/C604S) | pCAGGS_ss_His_TEV_hCHR2* (591) | FreeStyle™ 293-F | X-ray crystallography (hLeptin:hLEP-$R_{CRH2}$ assembly), SEC-MALS |
| Human Leptin wt (22-167) | pTwist_ss_His_TEV_hLeptin (539/556) | FreeStyle™ 293-F | X-ray crystallography (hLeptin:hLEP-$R_{CRH2}$ assembly), SEC-MALS (Ext. Data Fig. 2c, 8a), SDS-PAGE (Ext. Data Fig. 2d), Cryo-EM (hLeptin:hLEP-$R_{ECD\text{-}tGCN4}$ assembly) |
| Mouse LEP-$R_{IgCRH2}$ (328-633; C602S) | pCAGGS_ss_His_TEV_mIgCHR2* (582) | FreeStyle™ 293-F | X-ray crystallography (mLeptin:mLEP-$R_{IgCRH2}$ assembly), SEC-MALS (Ext. Data Fig. 2c) |
| Mouse Leptin wt (21-167) | pET11d_mLeptin (681) | *E. coli* (refolding) | X-ray crystallography (mLeptin:mLEP-$R_{IgCRH2}$ assembly), SEC-MALS (Ext. Data Fig. 6b,d,g,h), BLI (Ext. Data Fig. 1c), Cryo-EM (mLeptin:mLEP-RECD-ΔFNII-tGCN4 assembly; for initial templates in Ext. Data Fig. 8b) |
| Human LEP-$R_{IgCRH2}$ (330-635; C604S) | pCAGGS_ss_His_TEV_hIgCHR2* (581) | FreeStyle™ 293-F | SEC-MALS (Ext. Data Fig. 2c), SDS-PAGE (Ext. Data Fig. 2d) |
| Mouse Leptin wt (22-167) | pET15b_His_TEV_mLEP_WT (774) | *E. coli* (refolding) | Cryo-EM and SAXS (mLeptin:mLEP-$R_{ECD}$ and mLeptin:mLEP-$R_{ECD\text{-}tGCN4}$ assemblies), SEC-MALS (Fig. 2a, Ext. Data Fig. 1b, Ext. Data Fig. 6d,e,l, Ext. Data Fig. 7a), AUC (Fig. 2b-c, Ext. Data Fig. 6a), Signaling (Fig. 2d, Ext. Data Fig. 6k) |
| Mouse LEP-$R_{CRH2}$ (426-633; C602S) | pCAGGS_ss_HIS_TEV_mCHR2* (569) | FreeStyle™ 293-F | BLI (Ext. Data Fig. 1c) |
| Mouse LEP-$R_{ECD}$ (22-839) | pCAGGS_ss_HIS_Caspase3site_mLepR (635) | FreeStyle™ 293-F | BLI (Ext. Data Fig. 1c), AUC (Ext. Data Fig. 6a), SEC-MALS (Fig. 2a, Ext. Data Fig. 6b,d,h,i), AUC (Fig. 2b-c, Ext. Data Fig. 6a), MS Disulfide mapping (Ext. Data Fig. 5f), Cryo-EM and SAXS (mLEP-$R_{ECD}$ and mLeptin:mLEP-$R_{ECD}$ assembly) |

| | | | |
|---|---|---|---|
| Human Leptin$_{a1}$ (22-167; S141A/T142A) | pTwist_ss_His_TEV_hLEPa1 (598) | FreeStyle™ 293-F | SEC-MALS (Ext. Data Fig. 2c) |
| Mouse Leptin$_{a1}$ (22-167; S141A/T142A) | pET15b_His_TEV_mLEPa1_S141A-T142A (775) | *E. coli* (refolding) | SEC-MALS (Fig. 2a), Signaling (Fig. 2d) |
| Mouse Leptin$_{a2}$ (22-167; 60LDFI63/AAAA) | pET15b_His_TEV_mLEPa2_LDFI/AAAA (776) | *E. coli* (refolding) | SEC-MALS (Fig. 2a), AUC (Fig. 2b), Signaling (Fig. 2d) |
| Mouse Leptin site I variant (22-167; Q155A/D156A/W159A) | pET15b_His_TEV_mLEPsiteI Q155A/D156A/W159A (777) | *E. coli* (refolding) | SEC-MALS (Fig. 2a), Signaling (Fig. 2d) |
| Human Leptin wt (22-167) | pET15b_His_TEV_hLEP_WT (787) | *E. coli* (refolding) | Signaling (Fig. 2d, Ext. Data Fig. 6k) |
| Mouse LEP-Rb (22-1162) | pSems_ss_ALFAtag_HAtag_3xGGS_mLEP-Rb (708) | HEK293T | Signaling (Fig. 2d) |
| Mouse LEP-R$_{ECD\text{-}A407E}$ (22-839; A407E) | pCAGGS_ss_HIS_Caspase3site_mLepR_A407E (739) | FreeStyle™ 293-F | SEC-MALS (Ext. Data Fig. 6b) |
| Mouse LEP-R$_{ECD\text{-}\Delta FNIII}$ (22-633; C602S) | pCAGGS_ss_HIS_Caspase3site_mIgCHR1IgCHR2* (632) | FreeStyle™ 293-F | SEC-MALS (Ext. Data Fig. 6c,e) |
| Human LEP-R$_{ECD}$ (22-839) | pCAGGS_ss_HIS_Caspase3site_hLepR (611) | FreeStyle™ 293-F | SEC-MALS (Ext. Data Fig. 6f,g,j) |
| Mibavademab Heavy Chain | pTwist_ss_11510H_mibavademab (797) | FreeStyle™ 293-F | SEC-MALS (Ext. Data Fig. 6j), Signaling (Ext. Data Fig. 6k) |
| Mibavademab Light Chain | pTwist_ss_11510L_mibavademab (798) | FreeStyle™ 293-F | SEC-MALS (Ext. Data Fig. 6j), Signaling (Ext. Data Fig. 6k) |
| Mouse LEP-R$_{ECD\text{-}\Delta FNIII}$ (22-633; C602S) | pCAGGS_ss_HIS_Caspase3site_mIgCHR1IgCHR2* (632) | FreeStyle™ 293-F | SEC-MALS (Ext. Data Fig. 6c) |
| Human Leptin wt (22-167) | pET20b_ss_His_Caspase3site_hLEP (725) | *E. coli* (secreted) | SEC-MALS (Ext. Data Fig. 6f,g,h) |

| | | | |
|---|---|---|---|
| Murinized Human Leptin (22-167; G139L) | pET15b_His_TEV_hLEP_G139L (786) | *E. coli* (refolding) | SEC-MALS (Ext. Data Fig. 6i) |
| "Murinized" Human Leptin (22-167; G139L/I85V/M68L/G132D) | pET15b_His_TEV_hLEP_G139L/I85V/M68L/G132D (800) | *E. coli* (refolding) | SEC-MALS (Ext. Data Fig. 6i) |
| "Murinized" Human Leptin mCD-loop (22-167; H118S/W121Q/A122T/E126Q/T127K/L128P/D129E) | pET15b_His_TEV_hLEP_mCDloop (801) | *E. coli* (refolding) | SEC-MALS (Ext. Data Fig. 6i) |
| Human LEP-Rb (22-1165) | pSems_ss_ALFAtag_HAtag_3xGGS_hLEP-Rb (707) | HEK293T | Signaling (Ext. Data Fig. 6k) |
| Human Leptin$_{a1}$ (22-167; S141A/T142A) | pET15b_His_TEV_hLEP_S141-T142A (788) | *E. coli* (refolding) | Signaling (Ext. Data Fig. 6k) |
| Human Leptin$_{a2}$ (22-167; 60LDFI63/AAAA) | pET15b_His_TEV_hLEP_60AAAA63 (789) | *E. coli* (refolding) | Signaling (Ext. Data Fig. 6k) |
| Human Leptin site I variant (22-167; Q155A/D156A/W159A) | pET15b_His_TEV_hLEPsiteI_Q155A_D156A_W159A (793) | *E. coli* (refolding) | Signaling (Ext. Data Fig. 6k) |
| Mouse LEP-R$_{ECD\text{-}tGCN4}$ (22-839) | pTwist_ss_mLEP-R_5xGGS_tGCN4 (715) | FreeStyle™ 293-F | Cryo-EM and SAXS (mLeptin:mLEP-R$_{ECD\text{-}tGCN4}$ assembly), SEC-MALS (Ext. Data Fig. 7a) |
| Human LEP-R$_{ECD\text{-}tGCN4}$ (22-839) | pTwist_ss_hLEP-R_5xGGS_tGCN4 (713) | FreeStyle™ 293-F | Cryo-EM (hLeptin:hLEP-R$_{ECD\text{-}tGCN4}$ assembly), SEC-MALS (Ext. Data Fig. 8a) |
| Mouse LEP-R$_{ECD\text{-}\Delta FNIII\text{-}tGCN4}$ (22-633; C602S) | pCAGGS_ss_HIS_Caspase3site_mIgCHR1IgCHR2*_5xGGS_tGCN4 (726) | FreeStyle™ 293-F | Cryo-EM (mLeptin:mLEP-R$_{ECD\text{-}\Delta FNII\text{-}tGCN4}$ assembly; for initial templates in Ext. Data Fig. 8b) |
| Mouse LEP-R$_{ECD\text{-}\Delta FNIII\text{-}tGCN4}$ (22-633; C602S) | pCAGGS_ss_HIS_Caspase3site_mIgCHR1IgCHR2*_5xGGS_tGCN4 (726) | FreeStyle™ 293-F | Cryo-EM (mLeptin:mLEP-R$_{ECD\text{-}\Delta FNII\text{-}tGCN4}$ assembly; for initial templates in Ext. Data Fig. 8b) |
| mLEP-Rb$_{\text{-}mXFP}$ (22-866, P399S) | pSems-leader-mXFP-mLEP-R (706) | HeLa | smTIRFM |

| | | | |
|---|---|---|---|
| mLEP-Rb$_{ALFA}$ (22-1165, P399S) | pSems-leader-ALFAtag-mLEP-Rb (708) | HeLa | smTIRFM |
| mLEP-Ra$_{ALFA}$ (22-896) | pSems-leader-ALFAtag-mLEP-Ra (858) | HeLa | smTIRFM |
| mLEP-R (22-866)-3xGGS-Foldon$_{ALFA}$ | pSems-leader-ALFAtag-mLEP-R (22-866)-3xGGS-Foldon (856) | HeLa | smTIRFM |
| hTpoR | pSems-leader-ALFAtag-hTpoR | HeLa | smTIRFM |
